# On the feasibility of temporal interference stimulation of human brains using two arrays of electrodes

**DOI:** 10.64898/2026.03.31.715653

**Authors:** Yu Huang

**Affiliations:** Soterix Medical Inc., Woodbridge, NJ 07095; Department of Computer Science, University of Colorado at Colorado Springs, Colorado Springs, CO 80918

## Abstract

**Background:** Conventional temporal interference stimulation (TI, TIS, or tTIS) leverages two pairs of electrodes to induce an interfering electrical field in the brain. Both computational and experimental studies show that TI can stimulate deep brain regions without significantly affecting shallow areas. While promising, optimization of the locations and dosages on these two pairs of electrodes for maximal focal modulation remains computationally challenging. We are the first to propose two arrays of electrodes instead of two or multiple pairs of electrodes to boost modulation focality. However, the optimization algorithm outputs too many electrodes with overlaps across two frequencies, making it difficult to implement in practice.

**Objective:** Based on recent progress in developing multi-channel TI devices and computational work on TI optimization, here we again advocate two-array TI with feasibility data.

**Methods & Results:** We give a review on major algorithms for TI optimization, and compare these algorithms over 25 individual heads across six brain targets. We show that the latest optimization algorithm for two-pair TI innately works for two-array TI with the fastest speed (under 30s) and a similar amount of electrodes as in multi-pair TI. At four of the six targets, this fastest algorithm for two-array TI achieves similar or even better focality than TI using up to 16 pairs of electrodes that takes days to optimize. We also show a hardware implementation of two-array TI using 10 electrodes on our 8-channel TI device. We argue that two-pair TI is only preferred when one does not care about modulation focality or when hardware is limited to only two current sources. We restate the focality-intensity tradeoff but in the context of TI and provide a first voxel-level map (at 4 mm resolution) of achievable focality and modulation strength by TI in the MNI-152 head template.

**Conclusions:** Compared to two-pair or multi-pair TI, we promote two-array TI for its similar performance in focality and lower cost in terms of both optimization time and electrodes needed. We hope this work will pave the way for future adoptions of two-array TI for more focal non-invasive deep brain stimulation.

## Introduction

Recently, transcranial temporal interference stimulation (TI, TIS, or tTIS) has shown promise as a non-invasive deep brain stimulation method (Brahma et al., 2025; Grossman et al., 2017; Violante et al., 2023). TI places two pairs of electrodes on the scalp surface and injects two alternating electrical currents of similar frequency (eg. 1000 and 1010 Hz). The induced electric field (E-field) from these two alternating currents interfere with each other and thus generate amplitude-modulated electrical stimulation containing both a fast oscillating carrier signal in the kHz range and a slowly oscillating modulation envelope in a frequency that exactly equals the difference frequency of the two signals (Huang and Parra, 2019). The premise of TI is that neurons only respond to the slower oscillation due to their property of low-pass filtering (Grossman et al., 2017). Most TI studies to date are based on the conventional setup that includes two pairs of electrodes (Grossman et al., 2017; Lee et al., 2020; Stoupis and Samaras, 2022; Violante et al., 2023), and will be named two-pair TI afterwards. To the best of our knowledge, very few studies can be found in the literature that expand TI to use multiple electrodes per frequency. We name this type of TI using an umbrella term, multi-channel TI, throughout this paper. A natural extension of the conventional two-pair TI is to use more than two pairs of electrodes. For example, four-pair TI could use two pairs of electrodes for each of the two stimulating frequencies (Lee et al., 2022). We will name this as multi-pair TI in this paper. Note that for multi-channel TI, electrodes do *not* have to come in pairs. Instead, they could form an array for each of the two stimulating currents. To the best of our knowledge, we are the first to propose this array setup for TI and have formulated a rigorous mathematical framework to optimize the modulated envelope (Huang et al., 2020). We call this two-array TI in this work. We limit the number of stimulating frequencies to be two as in the conventional TI setting, and will discuss recent work expanding this to more than two frequencies (Botzanowski et al., 2025; Zhu et al., 2019) in the Discussions section.

Similar to conventional high-definition (HD) transcranial electrical stimulation (TES) (Datta et al., 2009; Dmochowski et al., 2011), individual optimization of electrode placement for TI matters significantly to maximize stimulation focality at the brain target (Brahma et al., 2025). Due to its non-linearity and non-convexness (Huang et al., 2020), TI optimization is usually computationally intensive: it takes hours to compute for only one optimal solution for the two-array TI we previously proposed (Huang et al., 2020), and it takes even more time (days) to optimize for the two-pair and multi-pair TI when using exhaustive searching methods (Huang and Datta, 2021; Lee et al., 2022, 2020). Other search heuristics such as genetic algorithms (Stoupis and Samaras, 2022) are adopted but the computation time is not significantly reduced to allow for practical use in a clinical setting, and such heuristics output different local minima at different runs (Goldberg, 1989). To the best of our knowledge, there is only one published algorithm that linearizes the mathematical formulation of TI optimization and thus it successfully reduces computing time to less than a minute (Geng et al., 2025). However, the authors developed this fast algorithm for conventional two-pair TI only, without realizing its implications for two-array TI.

In this work, we leverage the published fast TI optimization algorithm (Geng et al., 2025) to study the feasibility of two-array TI for better stimulation focality. We first compare several algorithms, on 25 individual heads across six brain targets, for maximizing TI focality including exhaustive search for two-pair TI (Huang and Datta, 2021; Lee et al., 2020) and multi-pair TI (Lee et al., 2022), our previously published algorithm for two-array TI (Huang et al., 2020), and the fast TI algorithm (Geng et al., 2025) for two-pair and two-array TI. We provide a review on the mathematical details for each algorithm, as well as empirical results from heuristics such as genetic algorithms. We show that different two-pair montages solved by genetic algorithms can lead to similar focality at the target, and how the limited number of available electrodes affect the focality of two-array TI. We re-state the focality-modulation tradeoff in the context of TI to argue that different TI modalities (two-pair, two-array) and optimization criteria (max-modulation, max-focality) are simply different samples from the focality-modulation curve (or Pareto front, Wang et al., 2023), as this is usually not well acknowledged in the community of researchers applying TI technology. With the fast TI algorithm, we are able to query the maximal focality that can be achieved by TI at almost every single brain location, and compare this data to that from conventional HD-TES, giving us a first voxel-level (at 4 mm resolution) comparison between optimized TI vs HD-TES. Finally, we show implementation of two-array TI based on our 8-channel TI device. We hope that this work will pave the way for future clinical applications of multi-channel especially the two-array TI for more focused non-invasive deep brain stimulation.

## Materials and methods

### Simulation database

This work builds upon our previous study on inter-individual variability of TI stimulation that generated 25 individual head models (Brahma et al., 2025). Briefly, ten individual T1-weighted head magnetic resonance images (MRI) (ages 22–35) randomly selected from the 1200 Subjects Data Release in the Human Connectome Project (Van Essen et al., 2012) and fifteen subjects from the Neurodevelopment database (Richards and Xie, 2015) (ages 17–89) were fed into open-source TES modeling software ROAST (Huang et al., 2019). With a simple one-line command, ROAST generated individual lead-field data corresponding to the international 10-10 system with 72 electrodes (Klem et al., 1999), for each of these 25 subjects. To make sure proper calculation of focality defined below, we interpolated all lead field data from mesh nodes onto regular grids defined by the image voxels. These lead-field data were used for the subsequent studies comparing different TI optimization algorithms. For full modeling details, see Brahma et al., 2025. We also built a model for the MNI152 head (Grabner et al., 2006) following the same modeling pipeline and used this model for other studies such as benchmarking the algorithm speed and surveying achievable focality across the brain.

### Comparison of optimization algorithms maximizing TI focality

Throughout this work, focality is defined as the size of the brain volume that receives a modulation depth (MD) of at least half of that achieved at the target (Huang and Parra, 2019). To express this volume on an interpretable spatial scale, we take its cubic root, so that focality is reported in centimeters (cm) and can be read as the edge length of a cube of equivalent volume. A smaller value therefore indicates a more focal modulation. We implemented and compared these algorithms for maximizing TI focality (Fig. 1, A1): 1) exhaustive search (ES; Fig. 1, C1) for two-pair TI (Lee et al., 2020); 2) heuristics for two-pair TI, including genetic algorithm (GA; Fig. 1, C2, Stoupis and Samaras, 2022), surrogate model (SM, Wang and Shoemaker, 2014), simulated annealing (SA, Geman and Geman, 1984), and particle swarm optimization (PSO, Kennedy and Eberhart, 1995); 3) fast TI (fTI; Fig. 1, B2, B4, C4, D5) for two-pair (Geng et al., 2025); 4) exhaustive incremental search (EIS; Fig. 1, C1) for multi-pair TI (Lee et al., 2022); 5) sequential quadratic programming (SQP; Fig. 1, C3) for two-array TI (Huang et al., 2020); and 6) fTI for two-array (Fig. 1, B2, B4, C4, D4). The reader is referred to the Appendix and Table A1 there for a full review of mathematical formulations and implementation details for different TI algorithms. Briefly, to solve the objective function (Eq. 4, Table A1; Fig. 1, B1), ES/EIS use brute-force search to find the global minimum maximizing focality of two-pair and multi-pair TI; heuristic search such as GA offers a quick alternative to find the local minimum; SQP maximizes TI focality at the target based on solution from HD-TES (Eq. 1, Table A1); fTI converts the non-convex TI optimization (Eq. 4; Fig. 1, B1) into two convex optimization problems (Eq. 5; Fig. 1, B2) defined on the left and right part of the brain (Fig. 1, B4) and these two can be efficiently solved by the classic linear-constraint-minimal-variance (LCMV; Fig. 1, C4) algorithm (Dmochowski et al., 2011). Once solved, fTI matches the induced E-field (*E* and *E* in Eq. 5) at the target by linearly scaling the two solutions (*s*_*L*_ and *s*_*R*_, Eq. 5) such that the MD is maximized there (Fig. 1, C4). This is because only when the amplitudes of the two stimulating current sources are equal can the MD be maximized (i.e., *E*_*L*_ = *E*_*R*_, Huang and Parra, 2019).

**Fig. 1:**
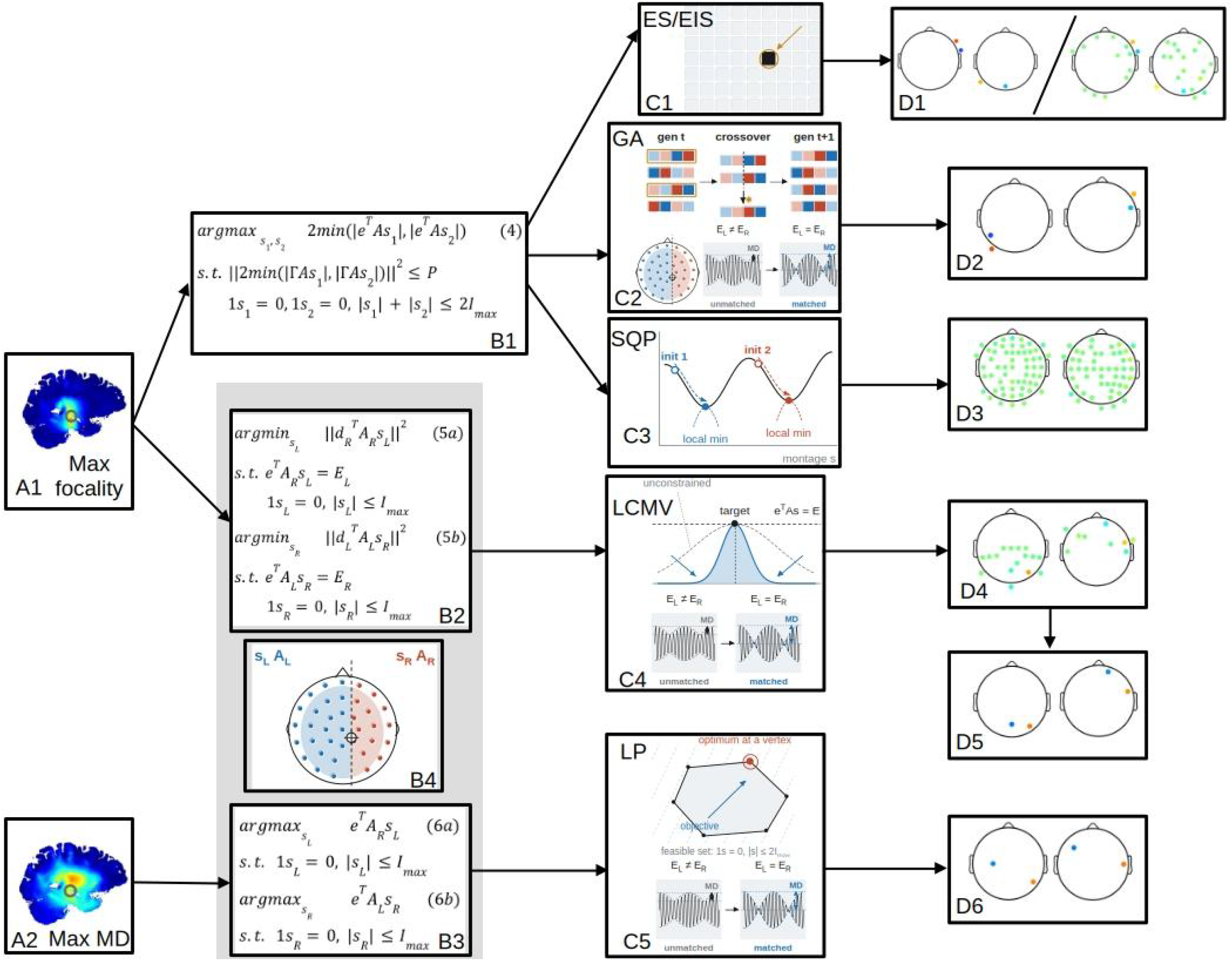
Overview of the optimization algorithms implemented in this work, for maximizing either TI focality (panel A1) or modulation depth (MD; panel A2). Taken from Table A1 in the Appendix, Eqs. 4 and 5 are the formulation for maximizing TI focality (panels B1, B2), and Eq. 6 is for max-MD (B3). Eqs. 5 and 6 are the fast TI formulation proposed by Geng et al., 2025 (indicated by the gray shaded area), and both require a split of the head model and candidate electrodes (B4). Eq. 4 can be solved by either exhaustive (incremental) search (ES/EIS; C1), heuristics such as genetic algorithm (GA; C2), or sequential quadratic programming (SQP; C3). As in Geng et al., 2025, Eqs. 5 and 6 can be solved by linear-constraint-minimal-variance (LCMV) and linear programming (LP), respectively, with a subsequent step of matching the electric field at the target (C4 & C5). This electric field matching, along with the split (B4), is also integrated in our implementation of heuristics such as GA (C2). ES and EIS give solutions for two pairs and multiple pairs of electrodes, respectively (D1). GA gives a two-pair solution (D2). SQP gives a two-array solution with many electrodes overlapping (D3). LCMV innately gives a two-array solution under the fast TI formulation for max-focality TI (D4), but Geng et al., 2025 chose the top two electrodes in terms of dosage for each frequency to obtain the two-pair solution (D5). Max-MD under the fast TI formulation solved by LP leads to a two-pair solution (D6). (Some of the panels were generated using Claude AI)

LCMV outputs array-like solutions (Fig. 1, D4), and fTI picks up the 2 electrodes in each frequency with top dosages to convert these array solutions into two pairs of electrodes (Fig. 1, D5). This greedy choice in fact compromises the stimulation focality, as we will see in the Results section. However, linearizing Eq. 4 to Eq. 5 significantly speeds up the computation (Table 1). We also adopted a similar idea in implementing all heuristic search algorithms such as GA for two-pair TI (Fig. 1, C2), i.e., assuming the two pairs of electrodes have the optimal locations on the left and right scalp, respectively, and matching the induced E-field at the target from the two pairs. This way the size of search space is greatly reduced. When solving Eq. 4 using SQP, we varied *P* for 12 different levels (Huang et al., 2020), each time initializing the TI optimization with solutions from HD-TES optimization; when solving Eq. 5 using fTI for both two-pair and two-array TI, we assigned *E*_*L*_ and *E*_*R*_ a series of 32 values from 0.01 V/m to 2 V/m. These parameters generate the focality-modulation curve of TI stimulation at the target (or Pareto front, Wang et al., 2023).

**Table 1:**
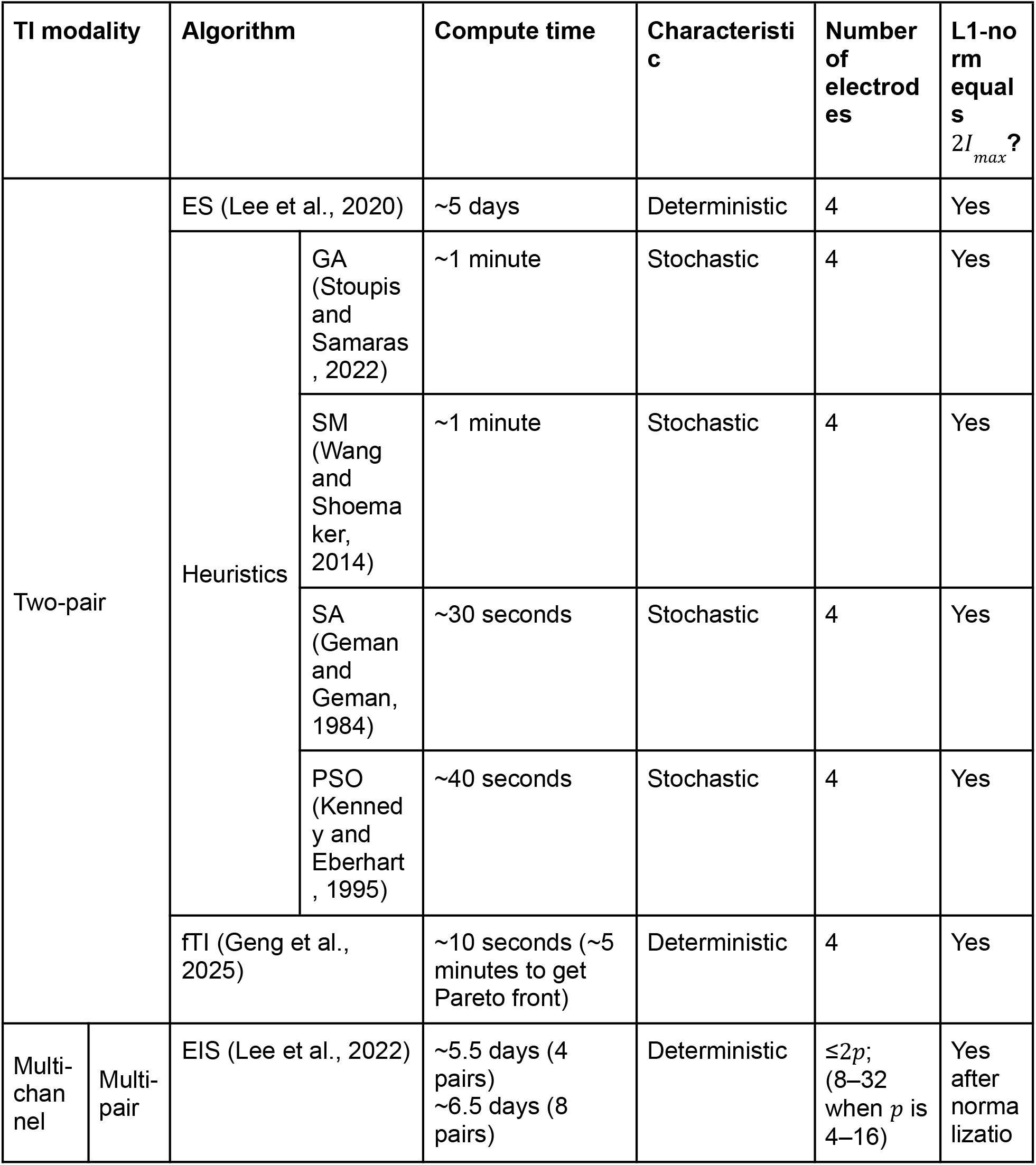
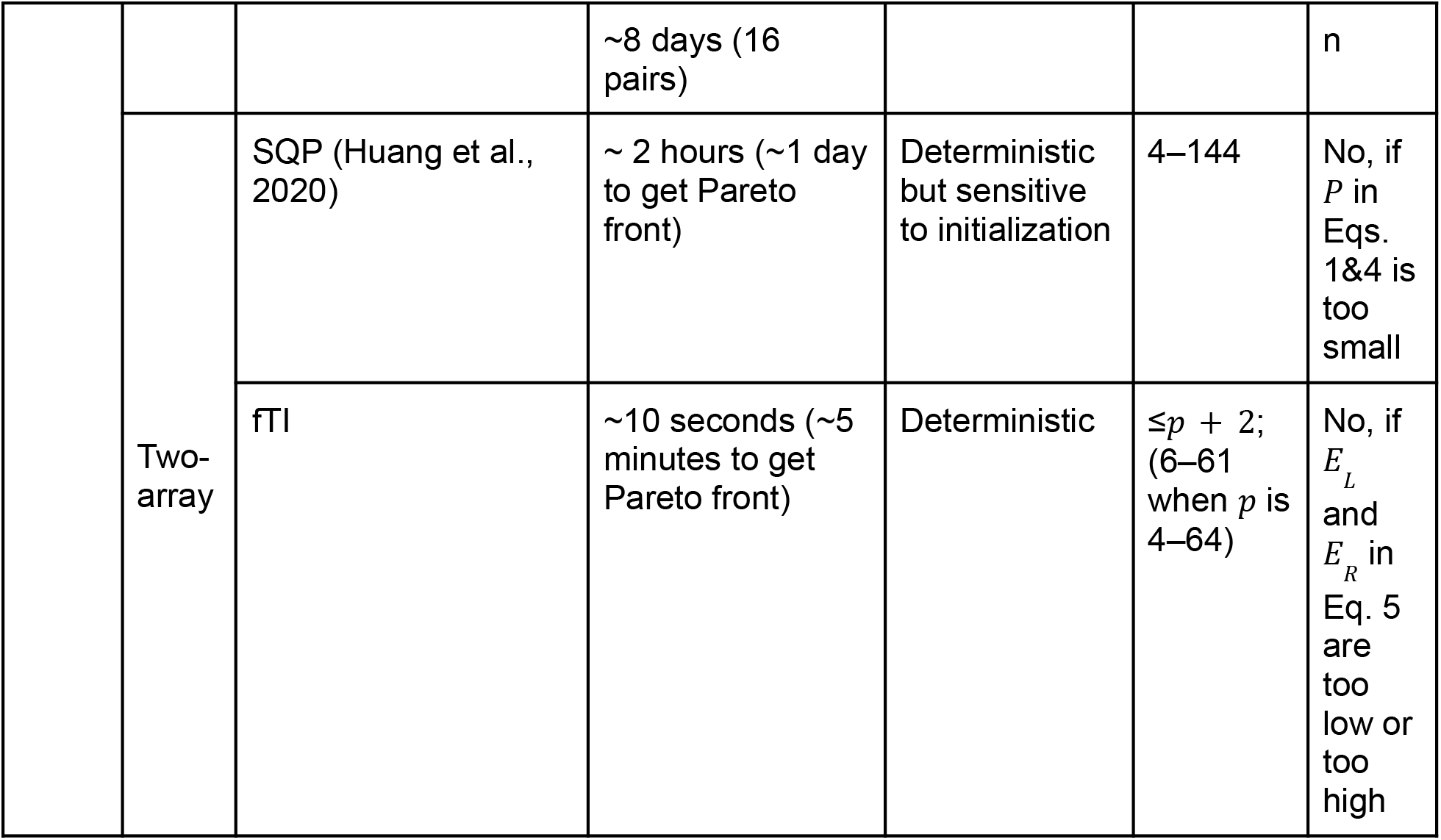
All algorithms for maximizing focality of TI compared in this work, with their respective compute time, specific characteristics, the number of electrodes needed, and the total current dosage (computed as the L1-norm of the solution vector) in the optimal montage. The compute time was the time it took to optimize for one target location from running Matlab R2022b on an Intel® Xeon(R) Silver 4216 CPU with 64 cores, and 128 GB RAM. *p* is the number of pairs in the multi-channel TI device. ES: exhaustive search; GA: genetic algorithm; SM: surrogate model; SA: simulated annealing; PSO: particle swarm optimization; EIS: exhaustive incremental search; SQP: sequential quadratic programming; fTI: fast TI algorithm.

We ran these algorithms on all the 25 individual head models described in the section of Simulation database, maximizing modulation focality in the radial-in direction at the same six locations in the brain as our previous work (Brahma et al., 2025): right hippocampus (MNI coordinates [28,-22,-14]), left dorsolateral prefrontal cortex (DLPFC, MNI coordinates [−39,34,37]), left motor cortex (MNI coordinates [−48,-8,50]), right amygdala (MNI coordinates [21,-1,-22]), right caudate (MNI coordinates [14,13,11]), and left thalamus (MNI coordinates [−9,-17,6]). Radial-in is the direction pointing from brain locations to the center of the brain defined as MNI coordinates [0,0,0]. We then compared the performance of these algorithms by looking at the achieved MD and its focality at the targets. For presentation, we first show these metrics in the focality-modulation space on Subject 1, and then show the averaged metrics with standard deviations on all the 25 subjects. To not clutter the figure, we show the results of two-pair TI and multi-channel (multi-pair and two-array) TI separately, and for two-pair TI, Pareto front generated by fTI was reduced to the single solution with the best focality when showing for the 25 subjects. Two-pair TI found by ES is also shown in the figure for multi-pair TI to show the trend of adding more pairs by the EIS algorithm. For multi-channel TI, Pareto front generated by ES/EIS, SQP, and fTI spans for different subjects at different locations in the focality-modulation space, due to inter-individual variability (Brahma et al., 2025). To facilitate comparison across these algorithms, we mapped the Pareto front to a normalized modulation depth between 0 % and 100 % by linear interpolation, where 0 % and 100 % are defined, respectively, as 0 V/m and the maximal MD achieved across subjects and algorithms at a specific target. When showing the results of two-array TI, solutions that use fewer than 4 electrodes and/or solutions with a total injected current dosage (L1-norm) not equal to 2*I*_*max*_ (4 mA) were removed. To get the number of electrodes in the solutions, we counted all the electrodes in *s*_1_ and *s*_2_ that have an absolute dosage of over 10^−6^ mA. Table 1 summarizes the number of electrodes and L1-norm of all the solutions. We also show the distributions of MD in the brain of Subject 1 and the corresponding optimal montages of some example data points in the focality-modulation space.

We compared the speed of each algorithm in Table 1. The speed depends on various factors such as hardware, software, size of head model, and parameter settings for the algorithms. To show a general trend in speed, we ran the algorithms with parameters implemented as described in the Appendix, on a workstation equipped with Intel® Xeon(R) Silver 4216 CPU with 64 cores, 128 GB RAM, and Matlab R2022b (MathWorks, Natick, MA). To generate the speed data, we ran the algorithms targeting the right hippocampus (MNI coordinates [28,-22,-14]) on the MNI152 head (Grabner et al., 2006).

### Stochasticity of the genetic algorithm

To study how stochastic the heuristics are for maximizing focality of the two-pair TI, we ran GA for 1000 times, targeting the right hippocampus (MNI coordinates [28,-22,-14]) on Subject 1. Each run lasted for 10 generations. The achieved MD and its focality at the target was recorded for each run and later analyzed. Note we used many short runs rather than a few long ones because the purpose of this analysis was to characterize the variability of solutions from GA, not to find a single best montage. Note also that our search space is smaller than in (Stoupis and Samaras, 2022), as each pair of electrodes is restricted to one of the two electrode subsets and the current amplitudes are set by E-field matching rather than searched over (see Appendix); the algorithm therefore converges in fewer generations.

### Effect of limiting the number of electrodes in two-array TI

To be compatible with our multi-channel TI device, we also limited the number of electrodes available in the fTI solutions for Eq. 5 (Fig. 1, B2). Our multi-channel TI device has up to 8 independent channels of current source, designed for TI using up to 8 pairs of electrodes. This means that the device can support up to 4 current sources per frequency. As we show in the hardware section below, 4 current sources can be converted into an array setup of up to 5 electrodes (the 5th one serving as return). Therefore, for each frequency (*s*_*L*_ or *s*_*R*_), we limit the number of available electrodes (*K*) to be one of these: 33, 17, 9, 5, 3, corresponding to a multi-pair TI setup of 64, 32, 16, 8, and 4 pairs of electrodes. While we have not developed any device that supports more than 8 pairs of electrodes, we still ran the fTI algorithm for theoretical results and to compare with the solutions from EIS for 16-, 8-, and 4-pair TI. To implement the constraint on available electrodes, we first solved Eq. 5 with all the 72 candidate electrodes placed by ROAST on the scalp, then chose *K* electrodes in the solution *s*_*L*_ or *s*_*R*_ with the top *K* absolute values, took out the lead field data in *A*_*L*_ or *A*_*R*_ corresponding to these *K* electrodes, re-ran the LCMV algorithms on this subset of lead field, and assigned the solution back to *s*_*L*_ or *s*_*R*_ at the corresponding electrodes. *s*_*L*_ and *s*_*R*_ were linearly scaled as the final solutions to make sure their induced E-field at the target (*E*_*L*_ and *E*_*R*_) were the same. We studied how different choices of *K* affect the achieved focality at the target output by the fTI algorithm.

### Focality-modulation tradeoff in TI

As we previously showed (Brahma et al., 2025; Huang et al., 2020), maximizing TI focality intrinsically encodes a tradeoff between the focality and the modulation at the target, and this tradeoff could be modeled by the energy constraint *P* as shown in Eq. 4 (Fig. 1, B1), or be controlled by the desired MD at the target (*E*_*L*_ and *E*_*R*_ in Eq. 5; Fig. 1, B2). Aside from max-focality described in previous sections, max-MD aims to maximize the MD at the target without considering its focality (Fig. 1, A2). We previously proved that max-MD solution will fuse the two pairs of electrodes in TI into one pair (Eq. 3 in Table A1, Huang et al., 2020), but Geng et al., 2025 showed that one can still solve for the max-MD solution if keeping the two pairs of electrodes separated (Eq. 6; Fig. 1, B3, B4). As we will see in the Results section, the max-MD solution with two pairs of electrodes separated always gives weaker MD compared to our “fused-pair” solution. More importantly, all these optimization criteria (max-focality, max-MD) and TI modalities (two-pair, two-array) are just samples from the focality-modulation curve (or Pareto front). To demonstrate this, we show the transition from max-focality to max-MD, giving us different TI modalities including two-array, two-pair TI, and eventually fusing of two pairs of electrodes leading to the max-intensity HD-TES (MD equals intensity in HD-TES). To this end, on Subject 1, we also solved Eqs. 3 and 6 targeting the right hippocampus (MNI coordinates [28,-22,-14]) and showed the solutions along with those from solving Eq. 5 with different levels of *E*_*L*_ and *E*_*R*_ (see Section of Comparison of optimization algorithms maximizing TI focality).

### Voxel-level maps of focality and modulation achievable by TI

We previously showed that two-array TI gives more focal stimulation than HD-TES, but this comparison is only limited to several locations in the brain (Brahma et al., 2025; Huang et al., 2020). Now that we can obtain the optimal two-array TI montage in under a minute thanks to the fTI algorithm, we could systematically check if this is true across the entire brain. To this end, we ran the fTI algorithm across the entire brain in the MNI152 head, maximizing the focality (Eq. 5; Fig. 1, B2) at each brain location, with desired MD (*E*_*L*_ and *E*_*R*_) set to 0.1 V/m. The result was compared to the max-focality solution of HD-TES (Eq. 2, Table A1) where *E* was specified as 0.1 V/m as well. We were also interested in how intense the modulation can be at each brain location, and thus ran the fTI algorithm again but this time we maximized the MD (Eq. 6; Fig. 1, B3, B4), giving us two-pair TI for each brain location. The result was compared to the max-intensity solution of HD-TES (i.e., fused-pair solution from Eq. 3 in Table A1). In all optimization runs the maximal current dosage (*I*_*max*_) was set to 2 mA. To make the computation finish in a reasonable amount of time, the algorithm was run on a grid of the MNI152 brain that was downsampled by a factor of 4 (i.e., 4 mm resolution), making this survey done on 21,911 locations in the brain.

### Hardware implementation of two-array TI

We recently built a prototype of an 8-channel TI stimulator. Each channel is electronically isolated and composed of one current source and one return electrode (i.e., a pair of electrodes). Therefore, 8 channels can support up to 8 pairs of electrodes for TI. However, in array setup, electrodes do not come in pairs. Instead, several electrodes form an array for each stimulation frequency. To convert our 8-channel stimulator into an array setup, we also manufactured a “merger” to merge all the return electrodes into one. We call this “Array Junction Box”, as it enables us to use our device to implement two-array TI. As the 4 return electrodes for each of the two stimulating frequencies are merged into 1, with two “mergers”, we can implement an array setup on our 8-channel device for a total of 10 electrodes (5 for each frequency, i.e., *K* = 5). See figure in the Results section for the entire setup.

We used this setup to show the feasibility of two-array TI on a tank filled with saline. Specifically, 10 stimulation electrodes were evenly placed at the inner periphery of the tank, fully immersed in the saline at the same height, with 5 on the left side for Frequency 1 and 5 on the right side for Frequency 2. For each frequency, 4 of the 5 electrodes were directly connected to the 8-channel stimulator at the anode sockets on the channel dispenser. The 5th electrode was connected to the 8-channel stimulator at the cathode sockets on the channel dispenser, but via the Array Junction Box that merges the 4 returns into 1. The stimulator was powered by a laptop computer and controlled by our HD-SC™ software platform. For a two-array TI setup, we intentionally assigned different dosages (in mA) for the 8 channels (from #1 to #8): 0.7, 0.9, 1.1, 1.3, 1.4, 0.6, 0.8, 1.1. Frequency was set to 1000 Hz for Channels 1–4, and 1010 Hz for Channels 5–8. Dosages and frequencies were set arbitrarily just to show feasibility. No phase delay was set up between channels. During stimulation, a Rigol DHO804 digital oscilloscope (RIGOL USA, Portland, OR) was used to probe the waveform in an arbitrary location in the saline. We observed the interfering waveform and confirmed interference in the saline.

## Results

### Comparison of stimulation focality across TI optimization algorithms

Figs. 2 and 3 show the achieved MD and its focality at the six target locations, for two-pair TI and multi-channel (multi-pair and two-array) TI, respectively. For both figures, the upper half (panel label starting with 1) shows the results for Subject 1, and the lower half (panel 2) shows the results (mean and standard deviation) for all the 25 subjects.

**Fig. 2:**
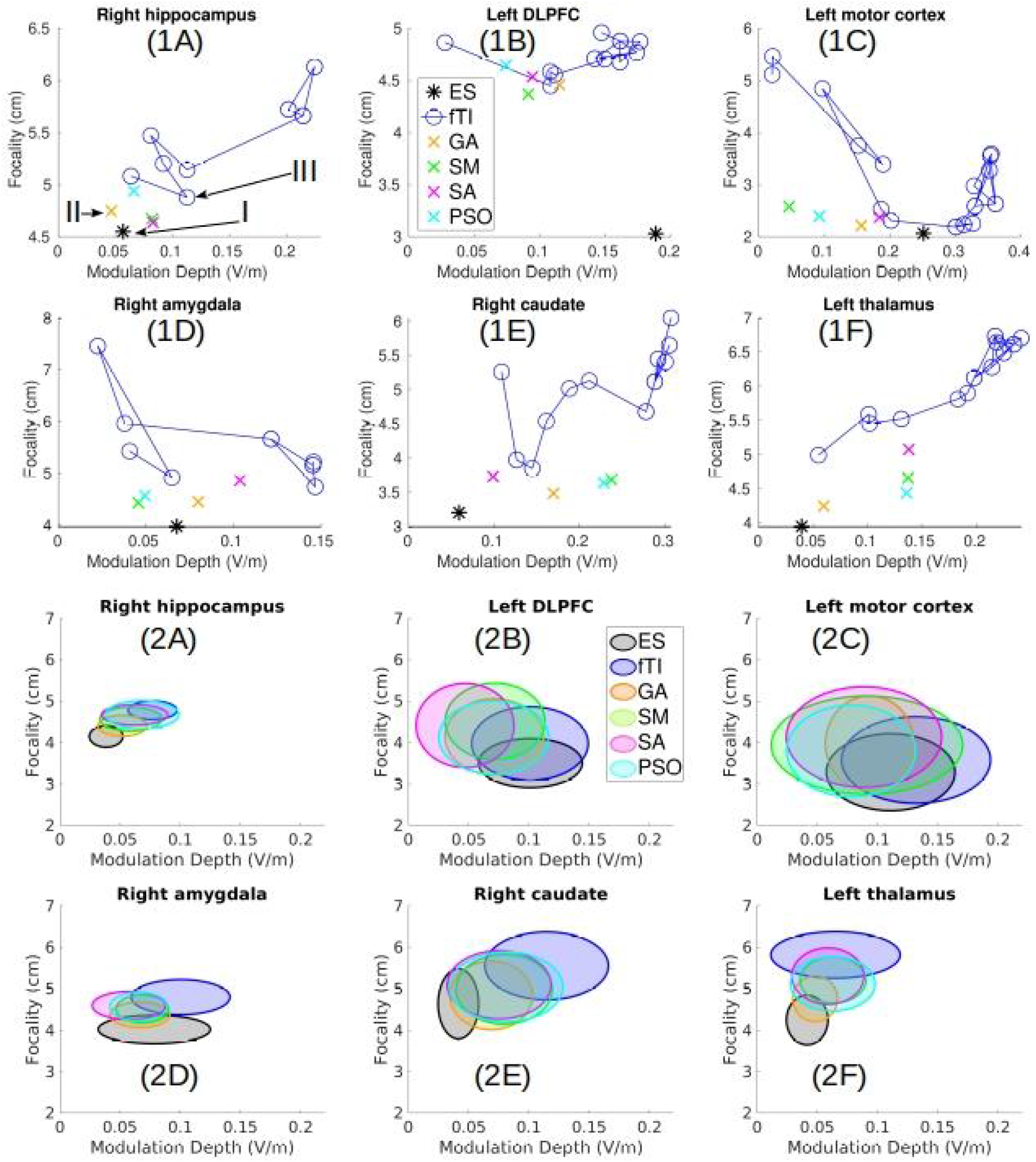
Focality-modulation plot of max-focality algorithms for two-pair TI, targeting six locations in the brain, for Subject 1 (panel 1), and for all the 25 subjects (panel 2). The center and spread of eclipse in panel 2 indicate the mean and standard deviation of the metrics (modulation depth and focality) across subjects along the respective axis. Note that the Pareto front of fTI shown in panel 1 (blue circles) is collapsed to the one with the best focality for each subject before averaging across subjects in panel 2. Arrows with Roman numerals in panel 1A indicate data points that are visualized in Fig. 4. ES: exhaustive search; GA: genetic algorithm; SM: surrogate model; SA: simulated annealing; PSO: particle swarm optimization; fTI: fast TI algorithm (for two-pair TI).

**Fig. 3:**
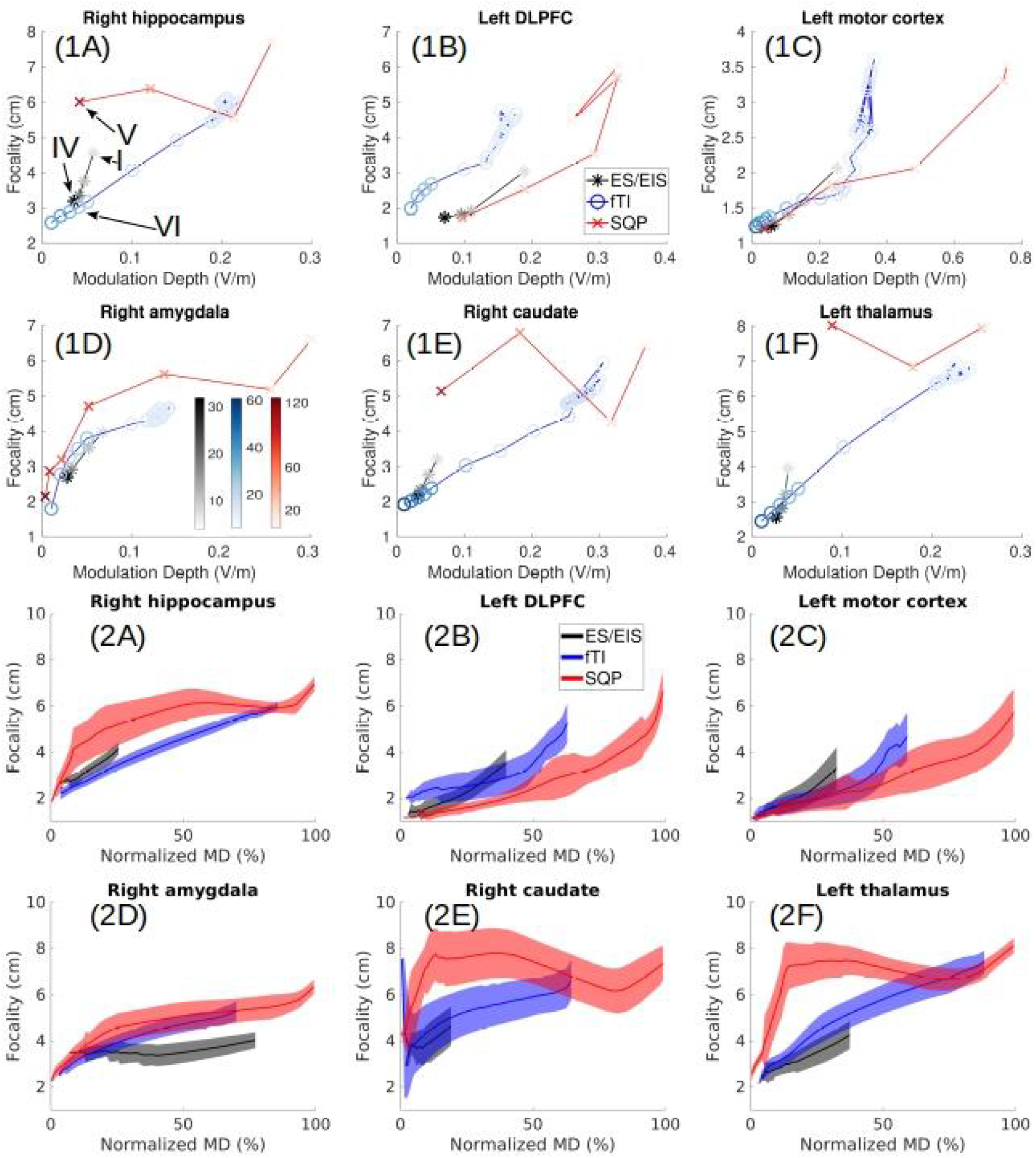
Focality-modulation plot of max-focality algorithms for multi-channel (multi-pair and two-array) TI, targeting six locations in the brain, for Subject 1 (panel 1), and for all the 25 subjects (panel 2). For pairs of electrodes, two-pair TI solutions are also plotted for reference (lightest gray asterisks). Colors on those symbols in panel 1 are coded by saturation (color bars in panel 1D) to indicate the number of electrodes in the optimized montage on Subject 1. The solid curve and shaded area in panel 2 indicate the mean and standard deviation in focality across subjects, with modulation depth (MD) normalized between 0 % and 100 %. Arrows with Roman numerals in panel 1A indicate data points that are visualized in Fig. 4. ES: exhaustive search (for two-pair TI); EIS: exhaustive incremental search (for multi-pair TI); SQP: sequential quadratic programming (for two-array TI); fTI: fast TI algorithm (for two-array TI).

For two-pair TI, it is clear that ES gives the best stimulation focality at all the targets for Subject 1 (black asterisks in Fig. 2, panel 1; best focality: 2.1 cm, panel 1C, shallow location), and also for all the subjects (center of black circles in Fig. 2, panel 2; best focality: ~3.3 cm in average, panel 2C), but it runs slowest (Table 1). fTI generates a Pareto front when it explores the search space for two-pair TI under different specified levels of MD at the target (*E*_*L*_, *E*_*R*_ in Eq. 5; Fig. 1, B2). The fact that it picks the top two electrodes in the solution of each frequency may contribute to the noisy and nonmonotonic property of the Pareto front (blue curve with circles in Fig. 2, panel 1). Note that even though fTI was run for 32 specified modulation levels (see Methods, Two-array TI), only part of these (8–16) runs are seen in Fig. 2, panel 1, as many of these runs converged to the same solution. When looking at the best solution in the Pareto front generated by fTI on Subject 1, it gives worse focality compared to ES and is on par with the heuristics (the lowest blue circles vs. asterisks and crosses in Fig. 2, panel 1), even though it runs fastest (but not the fastest in building the Pareto front; Table 1). When looking at all the subjects, fTI is significantly worse in focality compared to ES (blue vs. black, Fig. 2, panel 2; pair-wise t-test on focality with p value adjusted by Bonferroni correction: t(24)=26.1, p=10^−18^, panel 2A; t(24)=4.2, p=10^−3^, 2B; t(24)=4.4, p=10^−3^, 2C; t(24)=11.1, p=10^−10^, 2D; t(24)=10.8, p=10^−10^, 2E; t(24)=34.3, p=10^−21^, 2F), and the difference is bigger at deep locations (panels 2ADEF). The heuristics give significantly worse focality than that from ES (orange, green, magenta, cyan vs. black circles in Fig. 2, panel 2; Bonferroni corrected p<10^−3^ for all pair-wise t-tests), but they run much faster (Table 1). The heuristics are all significantly worse in focality than fTI at left motor cortex (orange, green, magenta, cyan vs. blue circles in Fig. 2, panels 2C; p<0.05 for all tests), but are significantly better than fTI at amygdala, caudate, and thalamus (panels 2DEF; p<0.05 for all tests). At the right hippocampus (Fig. 2A), GA and SM are significantly better in focality than fTI (p<0.05); at left DLPFC (Fig. 2B), SM and SA are significantly worse than fTI (p<0.05).

For multi-channel TI, EIS gets better focality when it finds more pairs of electrodes, for Subject 1 (black asterisks in Fig. 3, panel 1; best focality: ~1.2 cm, panel 1C; gray-level of blackness encodes number of electrodes in the solution with color scale shown in panel 1D; two-pair solutions are also plotted for reference, e.g., the asterisk labeled by Roman numeral I in panel 1A), and for all the subjects (black curves in Fig. 3, panel 2; best focality: ~1.2 cm in average, panel 2C). But EIS takes the most time to run (Table 1). Our old array optimization algorithm (SQP) performs on par with EIS in focality on Subject 1 for shallow targets (red crosses vs. black asterisks in Fig. 3, panels 1BC), but is worse for deep locations (panels 1ADEF). It also needs up to 120 (overlapping) electrodes (shown by redness levels of red crosses; color scale in panel 1D), and takes hours to run (Table 1). fTI generates more focal solutions than that from EIS on Subject 1, even more focal than 16-pair solutions, for all the deep target locations when using up to 61 electrodes (bluest circles in Fig. 3, panels 1ADEF; best focality: 1.8 cm, panel 1D). However, for shallow targets, fTI gives worse focality than that from EIS on Subject 1 (blue vs. black in Fig. 3, panels 1BC). When comparing these algorithms on all the subjects, at the normalized MD where different algorithms overlap, fTI gives significantly better focality than EIS for right hippocampus and left motor cortex (blue vs. black, Fig. 3, panels 2AC; two-way (modulation, method) repeated measures ANOVA: F(1,6)=74.1, p=10^−9^; F(1,16)=61.3, p=10^−8^), but is worse in focality compared to EIS at right amygdala (blue vs. black, panel 2D; F(1,29)=310.2, p=10^−15^) and left thalamus (panel 2F; F(1,19)=176.0, p=10^−12^), and is on par with EIS at left DLPFC (panel 2B; F(1,16)=1.3, p=0.27) and right caudate (panel 2E; F(1,7)=4.0, p=0.06). Interestingly, fTI performs significantly better in focality than our old algorithm (SQP) at all deep targets (blue vs. red, Fig. 3, panels 2ADEF; F(1,60)=84.2, p=10^−9^; F(1,55)=24.7, p=10^−5^; F(1,43)=240.2, p=10^−14^; F(1,51)=70.1, p=10^−8^), but is worse than SQP at shallow locations (panels 2BC; F(1,27)=116.7, p=10^−10^; F(1,38)=50.6, p=10^−7^).

To get a more intuitive view of how focal modulation each algorithm can achieve, we visualized six example solutions from panel 1A of Figs. 2&3, as indicated by the arrows with Roman numerals. Fig. 4 shows the electrode montages from these six examples and their corresponding distributions of MD. The electrode locations in the montage topoplots follow the international 10-10 convention (Klem et al., 1999). For two-pair TI, ES finds the most focal solution (4.55 cm focality, Fig. 4I) but it computes slowest (Table 1). fTI gives worse focality (4.88 cm, Fig. 4III) even though it’s much faster (Table 1). GA offers a nice tradeoff between speed and focality (4.75 cm, Fig. 4II), even though it is stochastic (Fig. 5). For multi-channel TI, 16-pair TI montage found by EIS gives better focality than two-pair solution (3.19 cm vs. 4.55 cm, Fig. 4IV vs. I). Our old algorithm (SQP) for array optimization does not converge on this subject at this target, which gives the worst focality (6.01 cm, Fig. 4V) and uses 108 electrodes (some of them overlap). fTI with array output generates the best solution with focality reaching up to 3.03 cm, even better than the 16-pair solution (Fig. 4VI vs. IV), but with much faster speed (Table 1) and a similar number of electrodes needed (25 in fTI vs 32 in 16-pair).

**Fig. 4:**
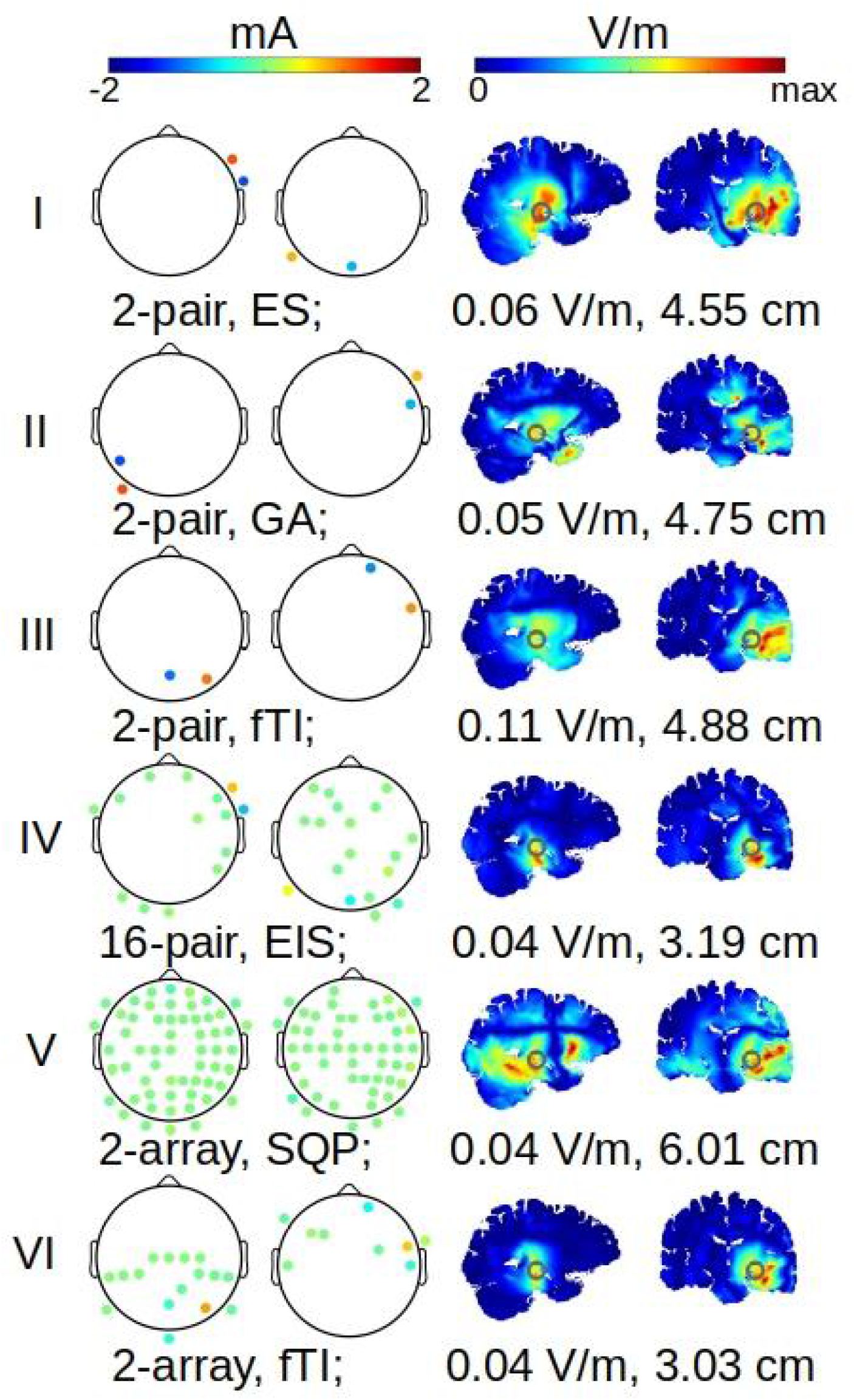
Visualization of data points taken out from panel 1A of Figs. 2 and 3. Roman numerals on the left link each row to the corresponding location in Figs. 2 and 3. The topoplots show the optimal electrode montages for the two frequencies used in TI on this subject (Subject 1), with electrode locations following international 10-10 convention. The heat maps show the achieved distribution of modulation depth (MD), in sagittal and coronal views, with target location (right hippocampus) shown as a gray circle. Color scales are shown on the top for the topoplots and heat maps, respectively. TI modality, algorithm name, MD and its focality at the target are noted below each row. ES: exhaustive search; GA: genetic algorithm; EIS: exhaustive incremental search; SQP: sequential quadratic programming; fTI: fast TI algorithm.

**Fig. 5:**
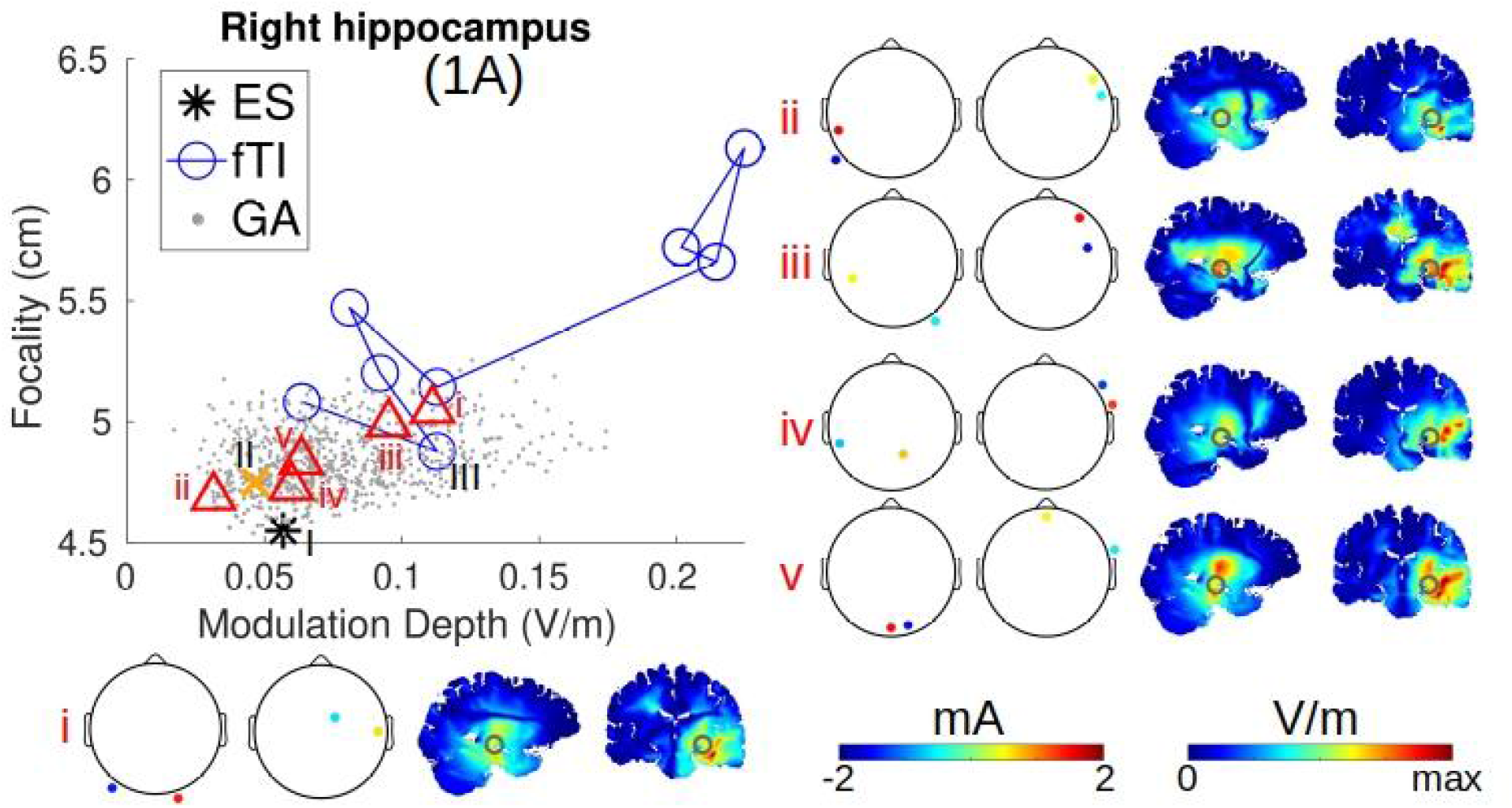
Stochasticity of the genetic algorithm (GA) for maximizing the focality of two-pair TI, shown across 1000 runs on Subject 1 targeting the right hippocampus. Gray dots in panel 1A are results from the 1000 GA runs. The first five runs are highlighted by red triangles labeled by red Roman numerals, and are visualized on the right for electrode montages (10-10 convention) and distributions of modulation depth (sagittal and coronal views). Target location (right hippocampus) is shown as a gray circle in the heat maps. Color scales are shown on the lower right for the topoplots and heat maps, respectively. The same data points of the exhaustive search (ES, black asterisk), GA (orange cross), and fast TI (fTI, blue circles) taken from Fig. 2, panel 1A are shown here in panel 1A for reference, with their visualization shown in Fig. 4 under Rows I, II, and III, respectively.

### Comparison of solution quality across TI optimization algorithms

It is clear from Table 1 that fTI is the fastest algorithm, with a single runtime only about 10 seconds (but takes ~5 minutes to build the Pareto front). Heuristics run a little slower but can still finish in 1 minute, but they are stochastic meaning one can get different results at different runs (Fig. 5). Our old array-optimization algorithm (SQP) finishes in 2 hours and takes a day to build the Pareto front. It is deterministic but very sensitive to initialization as we previously showed (Huang et al., 2020). ES takes days to finish but can get the deterministic global minimum. EIS takes one to three more days to find the multi-pair solution. All algorithms for two-pair optimization generate solutions with 4 electrodes and exactly matching the safety constraint encoded by the L1-norm of 2*I*_*max*_. EIS generates optimal montages with 8, 16, 32 electrodes for 4-, 8-, and 16-pair electrodes. Note that the number of electrodes can be smaller than 2*p* for *p* pairs of electrodes, as EIS could get stuck in local minima because if the focality is not improved when adding an extra pair of electrodes, the algorithm just keeps the existing solution (see Appendix for details). In our experiments searching for 4-, 8-, and 16-pair solutions on 25 heads across 6 targets, we found that only 40 of these 450 searches (8.9 %) got stuck in the local minima. Also one needs to normalize the final solution for EIS to match the safety constraint. Our old array algorithm SQP generates too many electrodes (up to 144) where some or all of the 72 electrodes overlap across the two stimulation frequencies. The algorithm also degrades to finding the two-pair solution when *P* is high (see Fig. 6 for more details on transition of TI modalities), leading to only 4 electrodes being used. Also when *P* is too small (strict), the algorithm gives a solution that cannot use up all the budget of current dosage (*I*_*max*_). fTI for two-array generates solutions with a reasonable number of electrodes (6–61) that is smaller than the number in the solution from SQP (up to 144). The number (*p* + 2) is also compatible with our multi-channel TI device supporting *p*-pair TI (see Hardware section). fTI for two-array gives solutions slightly violating the safety constraint (i.e., L1-norm slightly smaller or higher than 2*I*_*max*_), if *E*_*L*_ and *E*_*R*_ are specified too low or too high by the user.

**Fig. 6:**
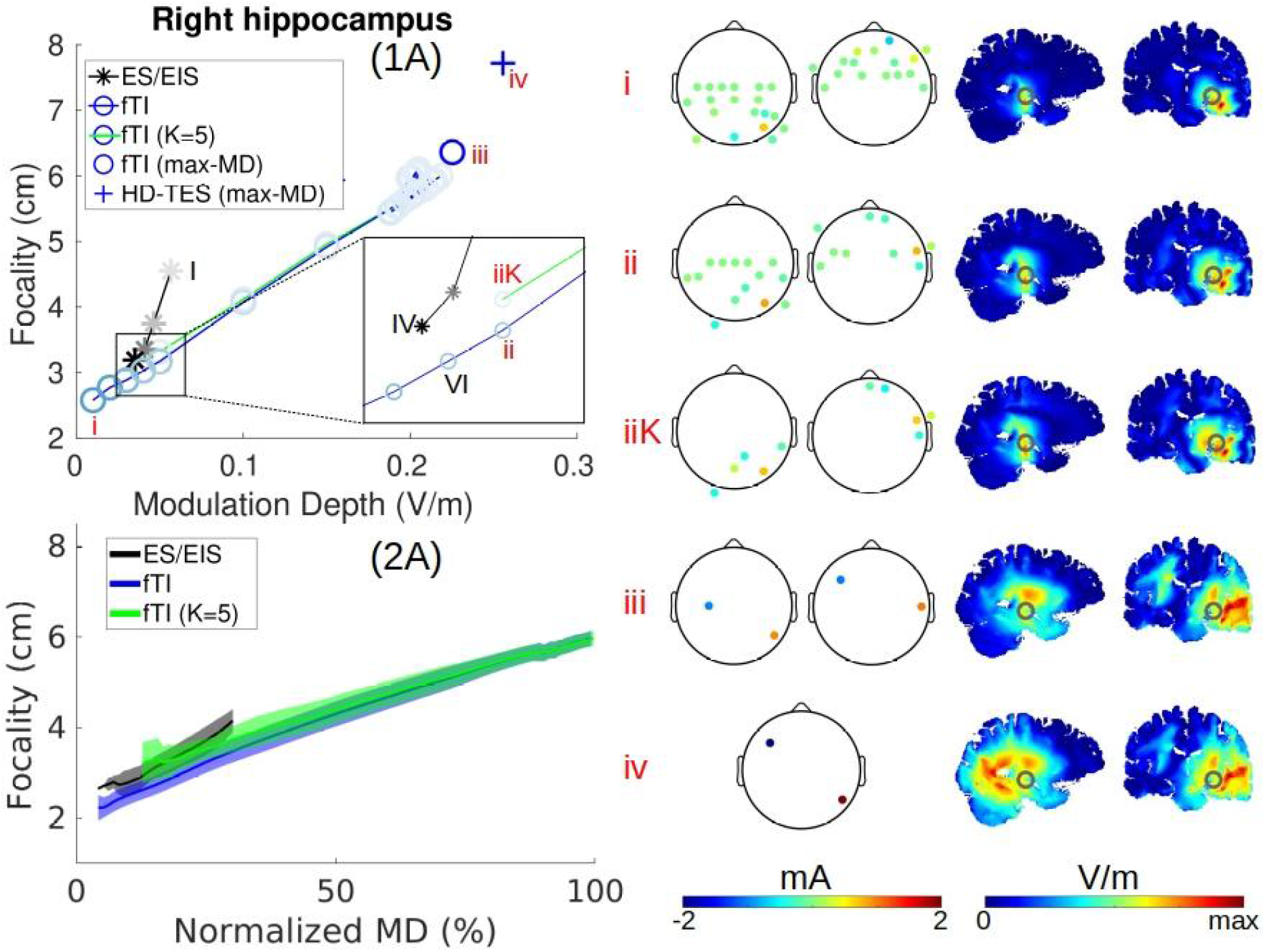
Effect of limiting number of electrodes for maximizing focality of two-array TI, and transition of max-focality two-array TI, to max-MD two-pair TI, and to max-MD HD-TES. Fast TI (fTI) algorithm maximizing focality for different levels of desired modulation at the target on Subject 1 is shown as a blue line with blue circles in panel 1A. fTI with *K* = 5 electrodes per frequency is shown as a green line with blue circles. Blue on the circles are coded by saturation (refer to Fig. 3, panel 1D for color bars) to indicate the number of electrodes in the solution. Inset in panel 1A shows the zoomed-in view of the data points. fTI maximizing modulation depth (MD) and our fused-pair (HD-TES) solution of maximizing MD are shown as blue circle (full saturation) and blue cross, respectively. Five data points labeled by red Roman numerals are selected from panel 1A for visualization on the right to show the transition of TI modalities (from i to iv) in electrode montages (10-10 convention) and MD distributions (sagittal and coronal views). Target location (right hippocampus) is shown as a gray circle in the heat maps. Color scales are shown on the lower right for the topoplots and heat maps, respectively. The same data points of the exhaustive (incremental) search (ES/EIS, black asterisk) and fTI taken from Fig. 3, panel 1A are shown here in panel 1A for reference, with their visualization shown in Fig. 4 under Rows I, IV, and VI, respectively. Panel 2A here shows the effect of limiting *K* = 5 in the fTI algorithm on all the 25 subjects targeting the same brain location (green curve and shaded area), with the same data of fTI without any limit on *K* (blue shaded area) and ES/EIS (black) from Fig. 3, panel 2A shown for reference.

### Stochasticity of the genetic algorithm

Fig. 5 panel 1A shows the distribution of the results from running the GA for 1000 times (gray dots) on Subject 1 targeting the right hippocampus. It is clear that the achieved focality is between those from fTI (blue circles) and ES (black asterisk), with the best focality being the same as that from ES (4.55 cm) and the worst reaching 5.26 cm. We also see that these 1000 runs can give us solutions reaching up to 0.17 V/m and down to 0.02 V/m in MD. Visualization of the first five runs (red triangles labeled by i–v) shows similar distributions of MD but with quite different optimal montages, highlighting the fact that the cost function of max-focality for two-pair TI has a highly non-linear and non-convex landscape. Therefore, many drastically different configurations of the two pairs of electrodes can lead to stimulation at the target with similar focality.

### Effect of limiting the number of available current sources in two-array TI

As the number of current sources (or channels) is a limited resource for two-array TI, we also studied how this affects the performance of fTI algorithm. Fig. 6 panel 1A shows this effect on Subject 1 targeting the right hippocampus. The green line with blue circles shows the result when limiting the number of current sources to be 4 per frequency (i.e., *K* = 5), corresponding to an 8-channel TI device (see Hardware section). It is clear from panel 1A that this constraint on *K* does not change the results much. It only compromises the focality by 0.15 cm compared to the one without this constraint (circles in the inset labeled by iiK and ii), while it greatly reduces the number of electrodes needed from 24 to only 10 that can be implemented on our device (topoplots in Fig. 6 labeled by ii and iiK; also Fig. 8). This 10-electrode TI solution performs almost the same compared to the 16-pair TI in focality (3.32 vs. 3.19 cm) that takes days to compute (iiK vs. IV in the inset of panel 1A, Fig. 6; the corresponding visualizations are in Fig. 6 and Fig. 4 labeled by these Roman numerals, respectively). For this subject and this target, other values of *K* (33, 17, 9) give exactly the same curve as the one without constraint (blue curve), and setting *K* to 3 makes the fTI algorithm not be able to return any meaningful solutions. Fig. 6 panel 2A shows the effect of limiting K for all the 25 subjects at this target, and the effect on all subjects at the other five targets is available in Supplementary Fig. S1. We can see that limiting *K* in the fTI algorithm only gives a small numerical difference in focality (green vs. blue, Fig. 6, panel 2A; Fig. S1). For our old algorithm SQP (Huang et al., 2020), we previously showed that limiting *K* gives solutions with worse focality (Huang and Datta, 2021).

### Focality-modulation tradeoff in TI

Fig. 6 shows the transition of TI modalities, obtained from optimization algorithms with different criteria. On the blue curve in panel 1A, we see transition from the one with most focal stimulation (i: focality 2.58 cm, MD 0.01 V/m, two-array TI with 35 electrodes), to the ones with intermediate focality and MD (VI: 3.03 cm, 0.04 V/m, two-array TI with 25 electrodes; ii: 3.17 cm, 0.05 V/m, two-array TI with 24 electrodes), to the one with most intense MD achievable under two-pair TI (iii: 6.36 cm, 0.23 V/m, two-pair TI with 4 electrodes), finally to the one with highest possible MD (iv: 7.72 cm, 0.26 V/m, fused-pair HD-TES with 2 electrodes). The visualization of these data points in Fig. 6 (i–iv) and Fig. 4VI clearly shows the transition from most focal to most intense modulation, and the need of more electrodes for more focal stimulation. This shows, under the context of TI, the focality-intensity tradeoff that was theoretically proved (Dmochowski et al., 2012). It also shows that it is possible to explore this tradeoff leveraging the LCMV algorithm embedded in the fTI algorithm to solve for the Pareto front (Geng et al., 2025; M. Wang et al., 2023).

### Voxel-level maps of focality and modulation achievable by TI

Fig. 7 shows the result of a survey we ran in the MNI152 head that sampled every 4 mm in the brain to query how focal or intense one can modulate using TI and HD-TES. Out of 21,911 brain locations we surveyed using the fTI algorithm, 21,834 (99.7%) returned meaningful results (LCMV did not converge on the others). We see that both HD-TES and TI can achieve focal stimulation at shallow cortical locations, but two-array TI excels in focal stimulation at deep brain regions (Fig. 7A, heat maps). A Wilcoxon signed-rank test shows that focality from two-array TI is significantly better than that from HD-TES (Fig. 7A, violins; medians: 3.27 cm vs. 4.06 cm for TI and HD-TES, respectively). The best focality achieved by two-array TI is 0.75 cm. Our previously proposed fused-pair solution for max-MD (Huang et al., 2020) indeed gives significantly higher modulation than that from two-pair TI computed by the max-MD fTI algorithm (Geng et al., 2025) (Fig. 7B, violins; medians: 0.28 V/m vs. 0.23 V/m for HD-TES and TI, respectively), especially at the shallow locations (Fig. 7B, heat maps). Under 2 mA stimulation, the MD can achieve up to 0.75 V/m in the brain by fused-pair HD-TES, consistent with our previous *in vivo* measurements (Huang et al., 2017).

**Fig. 7:**
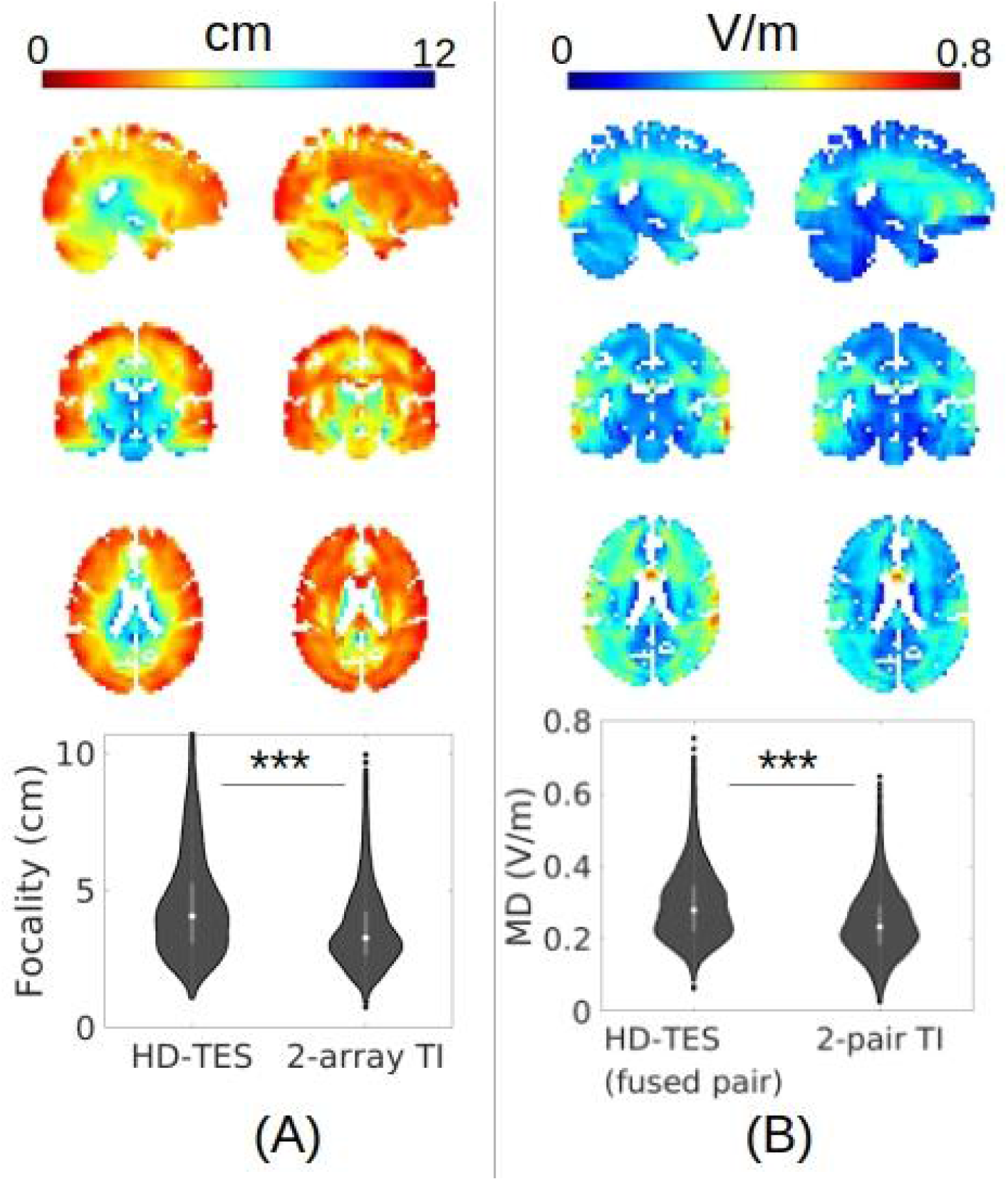
Voxel-level survey (at 4 mm resolution) on how focal (A) and intense (B) the modulation depth (MD) can be achieved at all the brain locations sampled at 4 mm in the MNI152 head, under a maximal current dosage (*I*_*max*_) of 2 mA. For panels A and B, the column on the left and right shows the voxel-wise distributions of metrics from HD-TES and TI, respectively. Heat maps are taken from the same brain slices. Color scales are shown on the top. The violin plots below show the metrics at all surveyed brain locations (N=21,834), with the median represented by the white dot, and the interquartile range indicated by the gray box. ***: p<0.001.

### Hardware implementation of two-array TI

Fig. 8 shows how we implement the two-array TI on our 8-channel TI stimulator, with the help of two Array Junction Boxes to merge the cathodes. Stimulation montage is shown in Fig. 8CD. 4 electrodes on the left side of the saline tank (a1–a4, Fig. 8CD) receive the currents of 1000 Hz from the first 4 channels of the stimulator (blue wires out of the channel dispenser, Fig. 8C), with the return electrode (cathode, c1, Fig. 8CD) going through the Array Junction Box and connecting back to the channel dispenser by 4 black wires (Fig. 8C, lower left). The other 5 electrodes stimulating at 1010 Hz on the right side of the saline tank follow a similar setup (Fig. 8A, upper right, and Fig. 8CD, right). During stimulation, we found strong interfering envelopes of 10 Hz at multiple locations in the saline (one example shown in Fig. 8E for 1 second period, i.e., 10 peaks), demonstrating the feasibility of two-array TI via our multi-channel TI hardware.

**Fig. 8:**
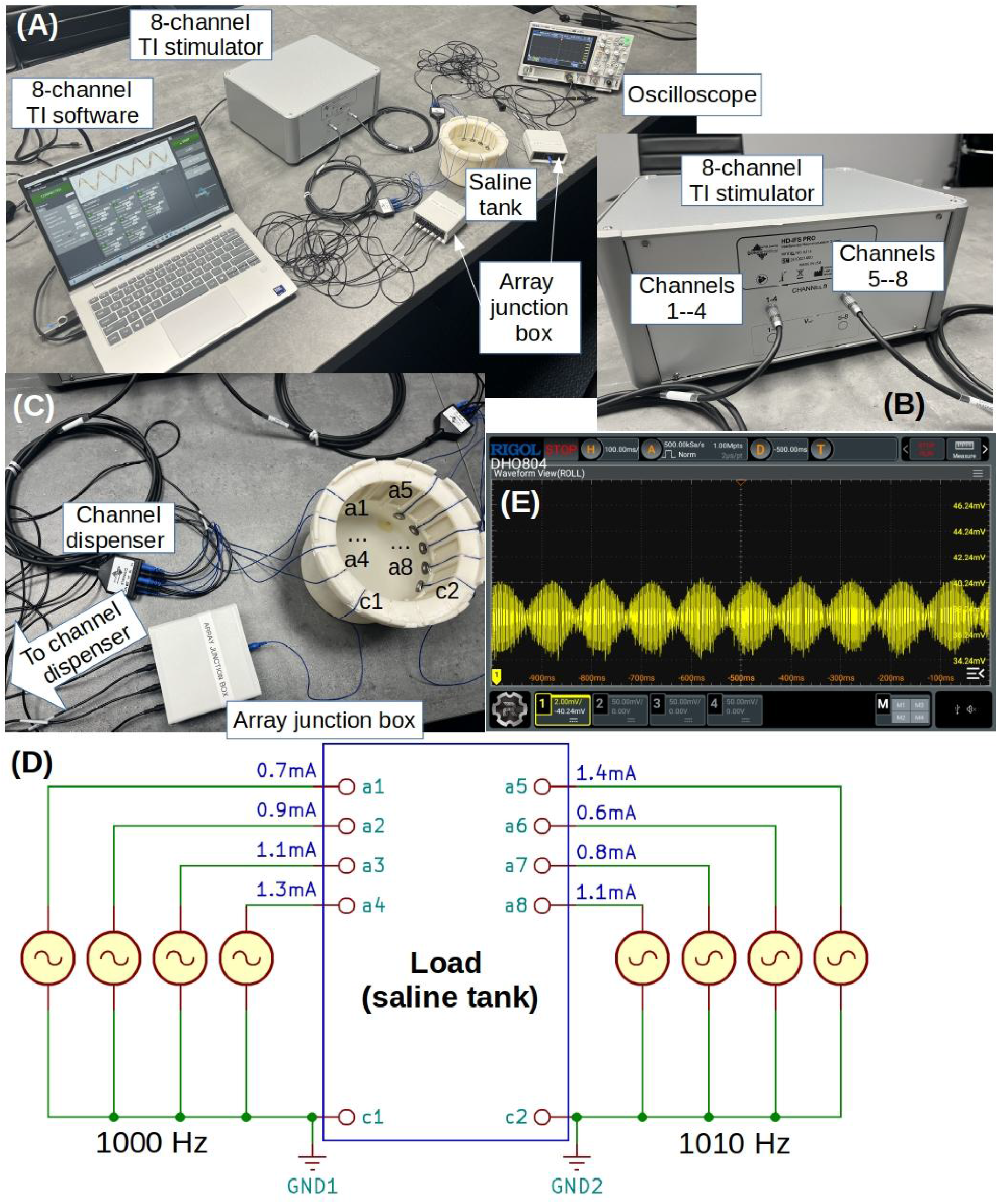
Implementation of two-array TI on our 8-channel TI stimulator. (A) Overall setup, with each major component labeled; (B) close view of our 8-channel TI stimulator, with channels labeled; (C) close view of the connections between stimulator and the saline tank for Channels 1–4, with Array Junction Box (“merger”) shown; electrodes on the saline tank are also labeled (a1 – a8: 8 anodes; c1–c2: 2 cathodes / return electrodes; corresponding to panel D); (D) circuit diagram of the 8-channel stimulator and the “merger”, with electrode names labeled and dosage for each channel shown; the anti-phase on the icons of current source on the left and right side of the load just means they are of different frequencies, but there is no phase delay between channels; (E) observed waveform in the saline for a duration of 1 second.

## Discussions

To the best of our knowledge, this work provides the first systematic review and evaluation of mainstream algorithms for TI optimization that covers two-pair, multi-pair, and two-array electrode setup. The evaluation was performed on 25 individual subjects across six brain targets including both shallow (cortical) and deep (subcortical) locations. We studied the performance of each algorithm in terms of the achieved focality, computation time, quality of the solution including number of electrodes needed and if it violates the safety constraint. We found that, for two-array TI, the fast TI algorithm (Geng et al., 2025) is the fastest and offers a similar and sometimes even higher focality than that from even 16-pair TI at some brain targets. For two-pair TI, heuristics such as the genetic algorithm (Stoupis and Samaras, 2022) offer a good tradeoff between speed and performance especially for deep targets, compared to the fast TI algorithm and exhaustive search. Overall TI with two arrays of electrodes can be more focal and faster to compute than using only two pairs of electrodes, even though a higher focality means a loss in modulation strength, an intrinsic tradeoff dictated by physics (Dmochowski et al., 2012). We further strengthen our work by showing that limiting the number of current sources in the optimal two-array TI montages only slightly reduces the focality, and by showing the implementation of a two-array TI using 10 electrodes on our 8-channel TI device.

The array optimization algorithm we published before (Huang et al., 2020) is shown to be sensitive to initialization, which may explain why in this work sometimes it performs even worse in focality than two-pair TI. More importantly, it outputs optimal montages with electrodes overlapping across the two interfering frequencies. This overlap introduces cross-talk across the two frequencies and will short-cut the current flow, resulting in almost no current flowing into the head. The fast TI algorithm avoids this problem by design, as it splits the candidate electrodes on the scalp into two halves in the first step, leading to an optimal montage that guarantees no overlap. Furthermore, as it is based on the classic LCMV algorithm, one can explicitly specify the desired modulation at the target as the linear constraint, which is more intuitive for practical implementations than our previous algorithm where one has to specify an abstract energy constraint (Huang et al., 2020). With the help of LCMV embedded in the fast TI algorithm, varying levels of desired modulation at the target naturally generates the focality-modulation curve, or the Pareto front when treating the TI optimization as a multi-objective problem (M. Wang et al., 2023). As we see in the Results section, TI with pairs of electrodes (both two pairs and multi pairs) has a Pareto front that lies in a different location in the focality-modulation space, and this Pareto front needs significantly more time to build than that from the fast TI algorithm that generates array-like solutions. This fact illustrates the complexity of the landscape of the cost function in TI optimization. We note that the contribution of this work is not the fast TI algorithm itself, which was developed by Geng et al., 2025, but the observation that this algorithm solves the two-array problem directly, and that the final step used to reduce its array solution to two pairs of electrodes is what limits the achievable focality.

We argue that the problem of maximizing TI focality with two pairs of electrodes is still not fully solved in practice. As a mixed-integer programming problem, it could be solved by the branch and bound (BnB) algorithm (Land and Doig, 1960). However, in our problem, we do not know if a better node (i.e., solution) hides under an unbounded node, so BnB degenerates to exhaustive search, which takes days to run. The fast TI algorithm innately generates array-like solutions, and Geng et al., 2025 chooses the top 2 electrodes in each frequency to convert the two arrays into two pairs. As we show in the Results section, this greedy choice in fact decreases the focality especially for the deep targets. Heuristics such as the genetic algorithm offer a good tradeoff between speed and performance. In fact, the NP-hardness of the mixed-integer programming (Papadimitriou, 1981) of maximizing the focality for two-pair TI, and the complex landscape of its non-convex cost function where many local minima exist, both make heuristics a nice choice, as exact algorithms will either easily get stuck in one of the many local minima or take much longer time to find the global minimum. Even though the genetic algorithm returns different montages at different runs, the spread in achieved focality across 1000 runs is modest (4.55 cm – 5.26 cm) as shown in the Results section.

We also argue that the problem of maximizing TI focality with more than two pairs of electrodes is still unsolved in practice. In this work, we used exhaustive incremental search to find multi-pair solutions based on the two-pair solution, as directly searching for, say, four pairs, would take significantly more time (11 billion combinations of electrodes to search through for four pairs vs. 1 million for two pairs). We also tried the genetic algorithm to search for a four-pair max-focality solution, but we found that it runs 3 times slower compared to two-pair under the same number of iterations, and the resulting focality is not necessarily improved over two-pair. On the other hand, the fast TI algorithm with array output can give us a solution in under a minute, with similar focality even when limiting the number of electrodes to match those in multi-pair setup. Nevertheless, developing fast and reliable algorithms for optimizing the focality of multi-pair TI remains an interesting challenge for future work. One could develop a highly parallelized version of heuristics to enable simultaneous explorations of the huge search space, or leverage recent progress in deep learning and AI (Bahn et al., 2022; Chen et al., 2025).

Based on the difficulty for finding optimal pair solutions mentioned above, we recommend against using pairs of electrodes for maximizing TI focality. Two-pair TI is only recommended for maximizing TI modulation without considering its focality. In fact, two-pair is just a special case in two-array setup where there is only 1 current source available for each frequency. Optimization of conventional HD-TES shows that max-intensity with safety (i.e., L1-norm) constraint always leads to montage with 1 current source (Dmochowski et al., 2011; Huang et al., 2020). Therefore, we argue that two-pair (1 source per frequency) is best for maximizing modulation of TI. For more focal TI, two-array offers significant advantages in both speed and focality. Note also that multi-pair TI is a restricted case of the array setup: requiring the electrodes to come in pairs constrains the montage vector to decompose into disjoint pairs of equal and opposite current, whereas in an array setup all electrodes of one frequency share a common return and the montage vector is constrained only by charge balance and by the safety limit on the total injected current. Two-array TI is therefore a generalization of multi-pair TI, and it is this larger feasible set that allows it to reach a more focal solution with fewer electrodes.

We note that even though the exhaustive search finds the global minimum that meets our max-focality objective function, the target location is not guaranteed to receive the maximal MD. This is due to the safety constraints we have imposed in the objective function. However, the target location still receives the most focal modulation found by the algorithm under this constraint. Achieving both focal and most intense MD under a fixed dosage budget is not possible (Dmochowski et al., 2012).

We provide a voxel-level map (at 4 mm resolution) of achievable focality and modulation in the brain by TI and HD-TES. Two-array TI gives significantly higher focality than HD-TES across the brain, but two-pair TI always outputs weaker modulation compared to HD-TES with one fused pair of electrodes. We learnt that in practice, one may want both focal and strong modulation in TI. While this is impossible per physics (Dmochowski et al., 2012), we offer some empirical guidelines: 1a) run max-focality algorithm to get an optimal montage for focal modulation; 1b) if the modulation is too weak, linearly scale up the dosage at each electrode; this will increase the modulation at the target without compromising the focality, but it may go beyond the safety limit posed on dosage; 2) alternatively, run max-MD algorithm to get the two-pair TI montage, or even better, run max-intensity for HD-TES to get the fused pair of electrodes if one does not care about the focality. Note that even if the fused-pair solution offers the strongest modulation, one may lose the perk of TI physics such as the rotational electric field (Huang, 2023; B. Wang et al., 2023) when fusing the two pairs of electrodes into one pair.

The purpose of the saline-tank experiment in this paper is only to show the feasibility of two-array TI, leveraging our 8-channel TI device and two Array Junction Boxes. Of course, the boxes could be integrated into the multi-channel TI device in the future, making a device exclusively for array-based TI. Note that the focality improvement shown in our models of two-array TI is still to be validated on either head phantoms or *in vivo* electrical recordings from human subjects.

In this work, we only focus on targeting a single location in the brain using TI with two interfering frequencies, and propose to use more electrodes to boost the modulation focality. Other works attempt to use more interfering frequencies to induce multi-focal TI stimulation or increase the single-focal TI focality. Specifically, Zhu et al., 2019 injects 4 sinusoidal signals with differing frequencies into two pairs of electrodes and was able to generate two focal spots of modulation, in both a head model and a petri dish filled with saline. However, they did not provide any mathematical framework on how to exactly steer the two focal spots especially in the head model, probably due to the increased complexity in physics of the 4 interfering signals. Botzanowski et al., 2025 showed that the modulation focality can be increased by adding more interfering frequencies, in recordings from both a mouse and a macaque monkey. But again no mathematical details are provided on how to control the location of the focal spot. We also note that Zhu et al., 2019 connects the power and ground of two of their 4 stimulating currents, essentially making 4 sinusoids into 2 amplitude-modulated signals, which is one of the cases shown in Botzanowski et al., 2025. Future work needs to figure out if it is possible to derive the analytic equations of the modulation depth related to the amplitudes of the multi (>2) interfering sinusoids, thereby enabling us to rigorously optimize and steer the current flow for multi-focal TI stimulation.

Several limitations of the underlying volume conductor modeling should be noted. All 25 individual head models, and the MNI152 model, were built from T1-weighted MRI alone. Segmentation accuracy, especially for the CSF and skull compartments, improves when a T2-weighted image is also available (Hoornweder et al., 2022), and errors in these compartments propagate to the predicted E-field and therefore to the MD and its focality. In addition, ROAST models the CSF as a single homogeneous compartment and does not explicitly represent the meninges; explicit emulation of the meningeal layers has been shown to change the magnitude and distribution of the predicted fields (Jiang et al., 2020; Weise et al., 2022). We note, however, that the present work is a comparison of optimization algorithms, and that all algorithms were run on exactly the same lead field data for each subject. Systematic biases in the forward model therefore affect all algorithms equally and are unlikely to change the relative ranking reported here, although the absolute values of the MD should be interpreted with this caveat in mind.

Finally, we caution that all the results reported in this work only look at the focality and strength of the low-frequency modulation, i.e., under the assumption that neurons only respond to the low-frequency envelope from the two interfering currents. We note that this mechanism of action still remains debated. For example, Mirzakhalili et al., 2020 shows that the TI stimulation waveform does not deliver any spectral energy at the difference frequencies (also known as “beat” frequency). Also, Opančar et al., 2025 and Semenov et al., 2025 show that the neurons and nerves also respond to high-frequency sinusoidal signals.

## Acknowledgement

The author would like to thank Abhishek Datta, Kamran Nazim, and Bhaskar Mukherjee at Soterix for insightful discussions on the hardware implementation of two-array TI. The author would also want to thank Kiril Balashov, Jonathan Romero, and the hardware team at Soterix for manufacturing the devices needed for experiments. The author would also like to thank the high-performance computing cluster at Boston University offered by Robert Reinhart as one of Soterix’s collaborators. This work was supported by the National Institute of Health through Grant R01DA060914, and by Soterix Medical Inc.

## Figure captions

**Fig. S1:**
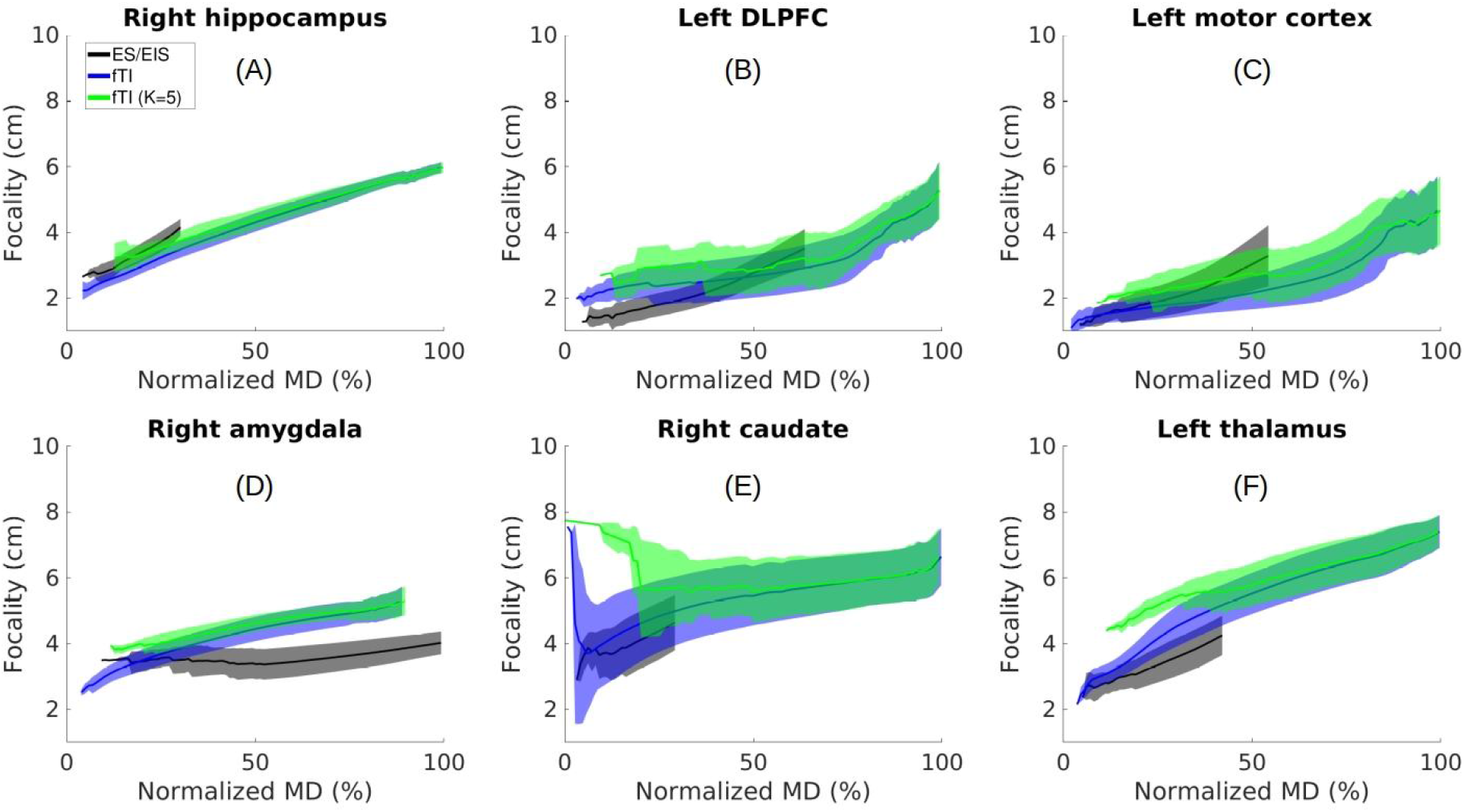
Same as Fig. 6 panel 2A in the main text, but for all the six targets studied in this work. Green curve and shaded area are results from limiting *K* = 5 in the fast TI (fTI) algorithm on all the 25 subjects, with the same data of fTI without any limit on *K* (blue shaded area) and exhaustive (incremental) search (ES/EIS, black) from Fig. 3, panel 2A shown for reference.

## Appendix Review of TI optimization algorithms

**Table A1:**
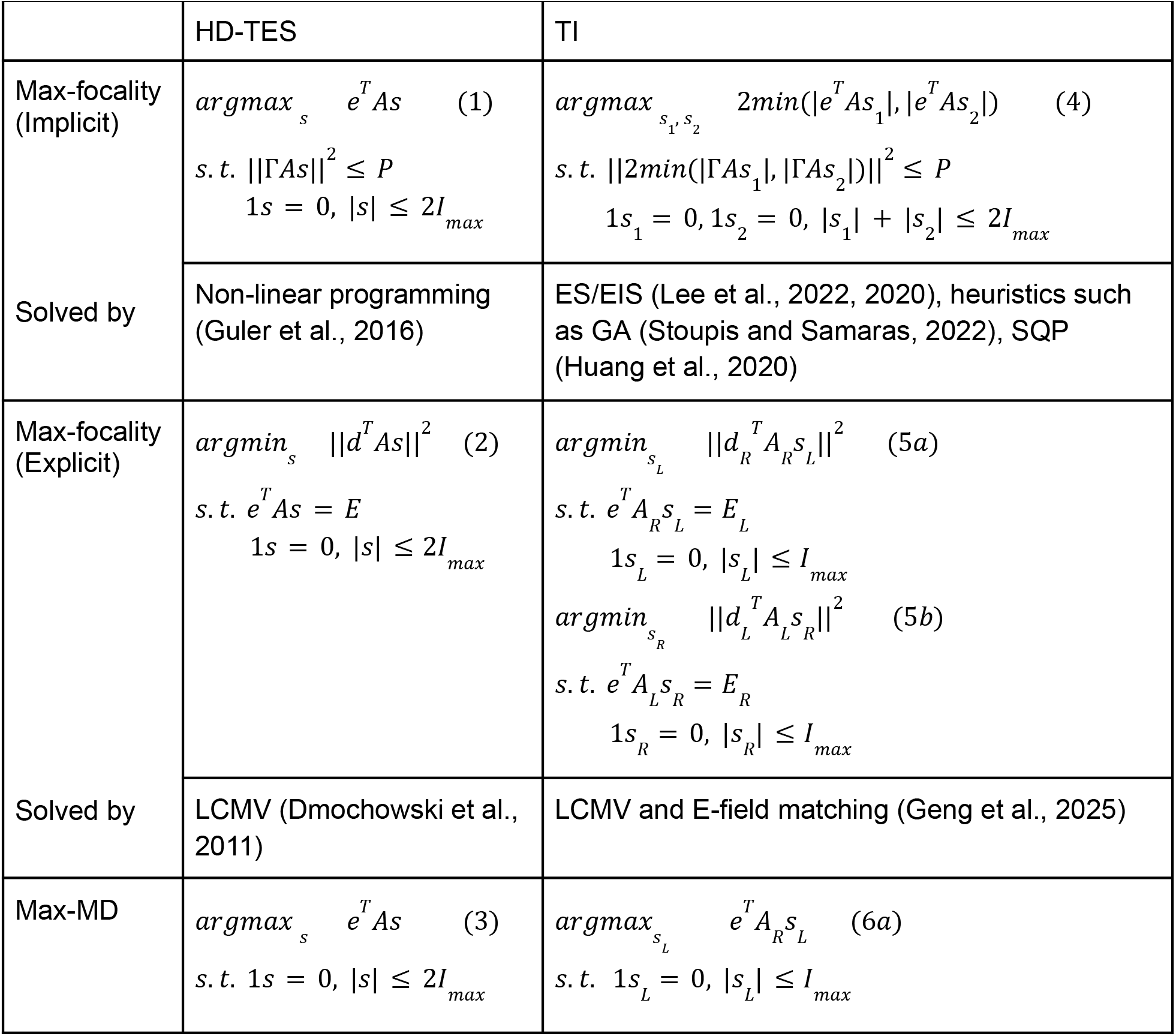
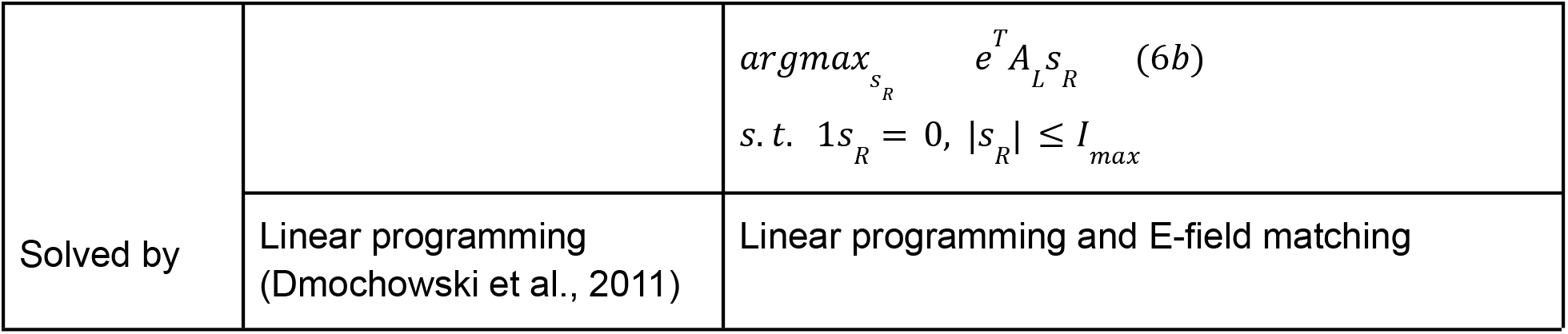
Summary of mathematical formulations for optimizing HD-TES and TI, with major algorithms for solving these equations noted. *A*: lead-field data generated from ROAST; Γ: spatial selectivity matrix that only selects the non-target locations; *s*: vector encoding the optimal electrode montage; *s*_1_, *s*_2_: optimal montages for the two stimulating frequencies in TI; *P*: user-specified energy threshold; *E*: desired electric field (E-field) at the target; *d*: orientations of the MD at all brain locations; *e*: orientation of the MD at the target (with values set to 0 at off-target locations); *d* and *e* are both set to radial-in throughout the paper; |. |: L1-norm; ||. ||: L2-norm; *T*: matrix transpose; *I*_*max*_: safety constraint on dosage; 1: a row vector with all values set to 1; subscripts *L* and *R*: the data in that notation corresponding to the left and right part of the brain (see text for details); MD: modulation depth; ES: exhaustive search; EIS: exhaustive incremental search; GA: genetic algorithm; SQP: sequential quadratic programming; LCMV: linear constraint minimal variance.

TI optimization builds upon optimization of the conventional HD-TES. In the most general mathematical formulation, the electric field (E-field, *e*^*T*^*AS* in Eq. 1) in HD-TES is replaced by the modulation depth in TI (MD, 2*min*(|*e* ^*T*^ *As*_1_ |, |*e*^*T*^ *As*_2_ |) in Eq. 4, Huang and Parra, 2019; Table A1). Note that the Max-focality criterion in Table A1 is divided into “implicit” and “explicit” formulations, as *P* in Eqs. 1&4 is an abstract quantity that is harder to specify than *E* in Eqs. 2&5. We will now walk through the major algorithms for solving TI optimization shown in Table A1.

### Max-focality

The goal of maximizing the focality of TI stimulation is simply to find the best montages of stimulation electrodes that can maximize MD at the target and meanwhile minimize the modulation energy at the off-target areas (Eq. 4). In the first constraint in Eq. 4, ||2*min*(|Γ*As*_1_|, |Γ*As*_2_|)||^2^ is the modulation energy at the off-target area (*min* and absolute operator || are done element-wise). The second constraint in Eq. 4 means that the montage vectors sum up to 0, and the total dosage (L1-norm of montage vectors) does not exceed a safety limit *I*_*max*_ (usually 2 mA). For more mathematical details, one is referred to Huang et al., 2020. Note that maximizing MD at the target and minimizing the modulation energy at the off-target areas are two contradictory objectives per physics (Dmochowski et al., 2012). Therefore, TI optimization formulated by Eq. 4 is a multi-objective optimization (M. Wang et al., 2023). As we see in the main text, solutions to the multi-objective optimization problem constitute a Pareto front.

As we can explicitly define the focality as the size of brain volume that receives at least half of the MD at the target (Huang and Parra, 2019), an alternative of Eq. 4 can be written as:

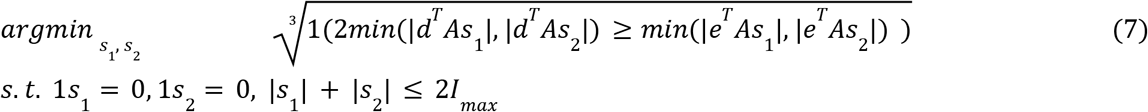

Here the MD at the target is 2*min*(|*e*^*T*^*As*_1_|, |*e*^*T*^ *As*_2_|), and MD at all the locations in the brain is 2*min*(|*d*^*T*^ *As*_1_|, |*d*^*T*^ *As*_2_|). The *min* and absolute operator || are performed element-wise. The inner product with the row vector 1 executes a sum representing the size of the volume. The cubic root brings this size to the 1D length scale. Also note that per our definition of focality in Eq. 7, a smaller number means more focal MD at the target, and thus we need to minimize.

#### Two-pair TI

A naive way of solving Eq. 7 is to search through all the candidate solutions, i.e., an exhaustive search (ES). We implemented the previously published method on searching (Huang and Datta, 2021; Lee et al., 2020). In details, the search was performed across: (1) four possible electrodes selected from the 72 placed electrodes on the scalp; (2) three possible ways of pairing in these four electrodes for the two frequencies; (3) amplitudes of injected current at one pair of electrodes, in the range of 0.5 mA–1.5 mA, with a step size of 0.05 mA and the sum across the two pairs of electrodes being 2 mA. In total 64,813,770 montages were searched for one target in the head model (1,028,790 possible combinations of 4 electrodes × 3 possible ways of pairing × 21 amplitudes of injected current). As we noted previously, this search takes days to complete.

For the two-pair TI, only four electrodes are used. The locations of these four electrodes could be modeled as integer variables to index the montage vectors. Therefore, for two-pair TI, Eq. 7 becomes a mixed-integer programming problem (Papadimitriou and Steiglitz, 1982). We can re-write Eq. 7 as:

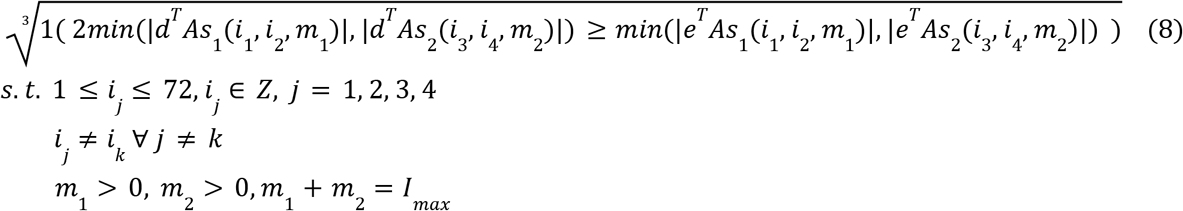

Here *i*_1_ to *i*_4_ are the integer variables representing the non-repeating indices of the four electrodes in montage vectors *s*_1_ and *s*_2_ (ranging from 1 to 72 as 72 electrodes were placed on the scalp by ROAST as candidates). *m*_1_ and *m*_2_ are positive non-integer values representing the current dosage for each of the two stimulating frequencies. The sum of *m*_1_ and *m*_2_ does not exceed the safety limit *I*_*max*_. We use the notation *s*_1_(*i*_1_, *i*_2_, *m*_1_) to indicate that *m*_1_ mA current goes into the *i*_1_-th electrode and flows out of the *i*_2_-th electrode in montage vector *s*_1_, with all other entries in *s*_1_ being set to 0. Similar meaning goes for *s*_2_(*i*_3_, *i*_4_, *m*_2_). As mixed-integer programming is NP-hard (Papadimitriou, 1981), one usually resorts to heuristic methods. One of the most commonly used heuristics in the TI literature is the genetic algorithm (GA, Stoupis and Samaras, 2022). Previous work using GA still takes hours to compute for only one target location. To reduce the size of the search space, we adopted a similar technique as in the fast TI (fTI) algorithm (Geng et al., 2025). Specifically, we divided the 72 candidate electrodes into two subsets (those to the left of the target location and those to the right, denoted by the subscripts *L* and *R* in Table A1), allowing one pair of electrodes to be chosen only from one of these subsets. Of course, this division could be done anterior-posterior, or superior-inferior, relative to the target location. We chose the direction that guarantees the most even split. This way we reduced the size of the search space from 72 to approximately 36 depending on the size of the subsets. We also did not explicitly search the non-integer variables *m*_1_ or *m*_2_. Instead, we linearly scaled the montage vectors inside the fitness function by matching the E-fields induced by *s*_1_ and *s*_2_ at the target (i.e., *E*_*L*_ and *E*_*R*_ in Eq. 5), a similar technique from fTI. This is based on the fact that the MD is maximized whenever the stimulations from the two frequencies are equal in strength (Huang and Parra, 2019). We used the function “gamultiobj” in Matlab (MathWorks, Natick, MA) to run GA, as “gamultiobj” can find the Pareto front. To this end, the fitness function implements the focality as shown in Eq. 8 and outputs both this focality and the MD at the target. The design vector consisted of four integer variables encoding the indices of the four stimulation electrodes, with the first pair constrained to the first subset and the second pair to the second subset of candidate electrodes; the current amplitudes were not encoded in the chromosome but were set analytically by matching the induced E-field at the target as just mentioned. We used the default settings of “gamultiobj”: a population size of 50, intermediate crossover with a crossover fraction of 0.8, adaptive feasible mutation, a Pareto fraction of 0.35 and crowding-distance based ranking. Bound constraints restricted each index to the range of the corresponding electrode subset. In the fitness function we penalized solutions that have electrode overlap or lead to an MD lower than 0.01 V/m at the target. Since GA is stochastic (Petrowski and Ben-Hamida, 2017), we ran “gamultiobj” six times and chose the solution with the best focality at each run. For speed, each run only lasted for 10 generations. We again chose the best focality within these six solutions as our solution from GA. The time for one run of “gamultiobj” is about 1 minute (Table 1 in main text).

We also implemented other heuristics as listed below:

1. Surrogate model (SM, Wang and Shoemaker, 2014). We used the Matlab function “surrogateopt” to build a smooth surrogate model to approximate the non-convex definition of focality in Eq. 8.
2. Simulated annealing (SA, Geman and Geman, 1984). We used the Matlab function “simulannealbnd”. We set a maximal number of 300 iterations for speed. As the landscape of the objective function is highly non-linear and non-convex, to make the algorithm perform more exploration rather than exploitation, we set the initial temperature to be high (10,000), and used a linear cooling schedule defined by the initial temperature subtracted by the iteration index. The algorithm was arbitrarily initialized by a montage of electrodes Fp2–F3 and O2–F7 for the two frequencies, respectively.
3. Particle swarm optimization (PSO, Kennedy and Eberhart, 1995). We used the Matlab function “particleswarm”. We set a maximal number of 3 iterations for speed. As the search space is huge, to make the algorithm explore more, we used a swarm size of 100 particles.

In these heuristics, the objective function was implemented the same way as in GA (except that only focality is output). The same techniques were used to reduce the size of search space, i.e., splitting the electrodes into two subsets, and matching the E-fields induced by *s*_1_ and *s*_2_ at the target. All these heuristics run under 1 minute (Table 1).

We also implemented the fTI algorithm for the two-pair TI (Geng et al., 2025). The key idea is to linearize non-convex Eq. 4 by splitting the head model and the electrodes into two subsets denoted by those notations with subscripts *L* and *R* in Eq. 5 in Table A1, maximizing the focality of stimulation for each subset using classic linear-constraint-minimal-variance (LCMV) algorithm for HD-TES optimization (Dmochowski et al., 2011), and matching the E-field at the target by linearly scaling the solutions for each subset. The LCMV algorithm gives an array of electrodes, instead of a pair of electrodes. Therefore, Geng et al., 2025 chose the top two electrodes in each subset solution before the matching. Here we implemented this for two-pair TI as well. Linearizing Eq. 4 significantly reduced the computation time (Table 1), but choosing the top two electrodes in the solutions is a greedy approach that compromised the focality, as we see in the Results section in the main text.

#### Multi-pair TI

For optimizing multi-pair TI, the goal is similar to Eq. 8, except that now we have 2*p* integer variables to encode electrode locations and *p* non-integer variables to encode current dosages (*p* is the number of pairs). One could still solve it using exhaustive search, but the size of search space increases exponentially with *p*. For example, when *p* = 2 (two-pair TI), there are 1,028,790 possible combinations of 4 out of 72 electrodes; when *p* = 4 (four-pair TI), this number jumps to over 11 billion. Therefore, an exhaustive search for multi-pair is computationally prohibitive unless one extensively parallelizes the search. Here, we implemented a heuristic based on the solution of two-pair TI, proposed by Lee et al., 2022. Starting from the exhaustive search for the optimal solution of two-pair TI at the same target, we searched for the next best pair of electrodes to be added to the solution. Specifically, once we had the optimal two-pair TI montage, we searched through the 3rd pair from the remaining 68 electrodes. For each pair candidate, we also searched through the current dosage in the range of −1 – 1 mA with a step size of 0.05 mA, and the options of adding this additional pair to *s*_1_ or *s*_2_. Once we have the three-pair solution, we searched for the 4th pair in the remaining 66 electrodes and repeated this process until the desired number of pairs was obtained. The final solution was normalized to make sure the total current injection does not exceed the safety limit (*I*_*max*_). We call this method exhaustive incremental search (EIS). For the comparison with other TI optimization algorithms, we searched for 4-, 8-, 16-pair solutions. It takes more time to search for more pairs, as shown in Table 1.

#### Two-array TI

As we proposed in Huang et al., 2020, electrodes do not have to come in pairs for multi-channel TI. Here we again implemented our algorithm for optimizing two-array TI to compare with other algorithms. Specifically, we first solved Eq. 1 for maximizing the focality of conventional HD-TES at the target location, using non-linear programming (Guler et al., 2016) implemented by Matlab function “fmincon”. We then used this solution to initialize the TI optimization (Eq. 4) and solved it by sequential quadratic programming (SQP, Brayton et al., 1979) implemented as the Matlab “fminimax” function. When solving, we varied *P* for 12 different levels (Huang et al., 2020), each time initializing the TI optimization with solutions from HD-TES optimization. This gives us the Pareto front of the optimization problem. This algorithm takes hours to solve for one target (Table 1).

To speed up the non-convex TI optimization, fTI algorithm (Geng et al., 2025) proposed a technique to linearize Eq. 4 by breaking it down into two HD-TES optimization problems, written equivalently in the form of LCMV (Eqs. 5a&5b) defined on the left and right part of the brain. As mentioned above, the split of the brain could also be done anterior-posterior or superior-inferior, depending on which direction generates the most even subsets. After Eq. 5 was solved, montage vectors *s*_*L*_ and *s*_*R*_ were linearly scaled to make sure their induced E-field at the target (*E*_*L*_ and *E*_*R*_) were the same. The scaled *s*_*L*_ and *s*_*R*_ were mapped back by their indices to the original montage vectors (*s*_1_ and *s*_2_ in Eq. 4) as the final solutions. Different from SQP for solving two-array TI where one has to specify an abstract energy constraint *P* in Eqs. 1&4, here one can specify a more intuitive *E*_*L*_ and *E*_*R*_ at the target as in Eq. 5 (and we always specified *E*_*L*_ and *E*_*R*_ to be the same value). One fTI run takes about only 10 seconds (Table 1). To generate the Pareto front using fTI algorithm for both two-pair and two-array TI, we assigned *E*_*L*_ and *E*_*R*_ a series of 32 values (in V/m): from 0.01 to 0.05 with step size of 0.01, to 0.9 with step 0.05, and finally to 2 with step 0.1. Note that LCMV will try to match *E*_*L*_ and *E*_*R*_ to this value initially, if not successful, it will reduce the value by 20% until the optimization converges. Therefore, the final solution does not necessarily guarantee the value one specifies. As LCMV generates an array of electrodes, this algorithm gives us two-array TI montages by default. Choosing the top two electrodes in the solution for each frequency will generate the two-pair solution, as originally proposed in Geng et al., 2025.

### Max-MD

We previously proved (Huang et al., 2020), both mathematically and numerically, that maximizing the MD induced by TI at the target will result in “fusing” of the two pairs of electrodes, with the locations of the “fused” pair of electrodes determined by solving the max-intensity problem of conventional HD-TES (Eq. 3 in Table A1). However, the fTI algorithm shows that one can still maximize the MD without fusing the two pairs of electrodes (Geng et al., 2025). Specifically, similar to Eq. 5, fTI decomposes the problem of maximizing MD of TI into two HD-TES max-intensity optimization as in Eqs. 6a&6b. After Eq. 6 was solved by linear programming, *s*_*L*_ and *s*_*R*_ were linearly scaled to make sure their induced E-field at the target (*E*_*L*_ and *E*_*R*_) were the same. As we see in the Results section in the main text, keeping the two pairs of electrodes separated (solution to Eq. 6) gives significantly weaker MD than using our previously proved fused-pair solution from Eq. 3.

